# *dotdotdot*: an automated approach to quantify multiplex single molecule fluorescent in situ hybridization (smFISH) images in complex tissues

**DOI:** 10.1101/781559

**Authors:** Kristen R. Maynard, Madhavi Tippani, Yoichiro Takahashi, BaDoi N. Phan, Thomas M. Hyde, Andrew E. Jaffe, Keri Martinowich

**Author notes:** Correspondence: Kristen R. Maynard, Lieber Institute for Brain Development, 855 North Wolfe Street, Suite 300, Baltimore, MD, 21205; Andrew E. Jaffe, Lieber Institute for Brain Development, 855 North Wolfe Street, Suite 336, Baltimore, MD, 21205.

## Abstract

Multiplex single-molecule fluorescent in situ hybridization (smFISH) is a powerful method for validating RNA sequencing and emerging spatial transcriptomic data, but quantification remains a computational challenge. We present a framework for generating and analyzing smFISH data in complex tissues while overcoming autofluorescence and increasing multiplexing capacity. We developed *dotdotdot* (https://github.com/LieberInstitute/dotdotdot) as a corresponding software package to quantify RNA transcripts in single nuclei and perform differential expression analysis. We first demonstrate robustness of our platform in single mouse neurons by quantifying differential expression of activity-regulated genes. We then quantify spatial gene expression in human dorsolateral prefrontal cortex (DLPFC) using spectral imaging and *dotdotdot* to mask lipofuscin autofluorescence. We lastly apply machine learning to predict cell types and perform downstream cell type-specific expression analysis. In summary, we provide experimental workflows, imaging acquisition and analytic strategies for quantification and biological interpretation of smFISH data in complex tissues.

## INTRODUCTION

In the age of rapidly advancing Next Generation Sequencing (NGS) technologies, such as single cell RNA-sequencing (RNA-seq) and spatial transcriptomics^1, 2^, single-molecule fluorescent *in situ* hybridization (smFISH) has emerged as a potential gold standard for validating and extending findings derived from large scale transcriptomic data. The widespread generation of single-cell RNA-seq data sets in the neurosciences has fueled a resurgence of smFISH approaches to validate cell type-specific molecular profiles by visualizing individual transcripts at cellular resolution^3, 4^. Information from single-cell RNA-seq data has revealed increasingly complex transcriptomic signatures for functionally distinct cell types, including the recently identified Rosehip neurons in cortical layer one^5^ and cells with neurogenic potential in the dentate gyrus of the hippocampus^6, 7^, such that molecular definition of these cells necessitates combinatorial labeling with multiple probes to confirm both presence and absence of specific transcripts within a spatially-defined context.

While chromogenic and fluorescent i*n situ* hybridization methodologies have been utilized for decades^8, 9^, recent advances in hybridization/probe technologies, imaging techniques, and data analysis tools have streamlined smFISH assays and improved sensitivity and specificity^10–12^. Despite these methodological advances, multiplexing in complex tissues with extensive cellular heterogeneity, such as postmortem human brain, remains a significant challenge. Studying human brain tissue is also complicated by high levels of autofluorescence due to lipofuscin granules^13, 14^. Indeed, to avoid confounding signals from lipofuscin, the majority of smFISH investigations in postmortem human brain have been limited to single or duplex chromogenic approaches^6, 7, 15–17^.

While studies have begun to incorporate multiplex fluorescent approaches in postmortem human brain tissue, no consistent strategy for eliminating, masking, or subtracting lipofuscin autofluorescence has been described^3, 5, 18–22^. Some studies have characterized lipofuscin autofluorescence based on size and intensity or custom filter cubes^3, 4, 21^, but these reports do not document how these approaches impact quantification of fluorescent signals from probe hybridization. An image processing approach in non-human primate brain tissue using spectral imaging and linear unmixing showed promise for characterizing and removing lipofuscin autofluorescence^23^. However, this approach has not yet been validated and widely implemented in postmortem human brain. Furthermore, the four dimensional data sets acquired using multispectral imaging across a tissue depth add additional computational hurdles for automating image analysis and quantifying single transcripts.

Several microscopy-based methodologies for single cell, spatially resolved transcriptomics have been developed^24–28^. However, these highly specialized platforms still rely on the availability and accuracy of algorithms for fluorescence segmentation, and often require sophisticated microscopy equipment and reagents that are not readily available to the majority of laboratories. Commercially available smFISH platforms have the capacity for higher-order multiplexing and can, in theory, be used for differential expression analysis within molecularly and spatially defined cell types. However, the downstream computational tools for analyzing these types of data have lagged behind their widespread use, and those tools that have been developed remain largely inaccessible to most neurobiology labs without strong computational expertise. Hence, the majority of current smFISH applications have been qualitative, rather than quantitative, and have therefore not maximized the utility of these potentially rich imaging datasets.

To address this need, we developed an intuitive and adaptable computational workflow called *dotdotdot* to quantify individual RNA transcripts at single cell resolution in intact tissues and performed differential expression analysis of smFISH data. We validate the accuracy of *dotdotdot* for quantifying RNA transcripts in both mouse and postmortem human brain and use computational approaches, such as K-means clustering and machine learning, to answer biological questions about gene co-expression and molecular cell type based on quantitative analysis of spatial gene expression. In summary, we present an imaging platform coupled with computational tools for smFISH data that can be readily implemented in most laboratories without need for highly specialized expertise or equipment to elevate spatial analyses of gene expression and complement growing single cell and spatial transcriptomic data sets in the field of neuroscience and beyond.

## RESULTS

### The *dotdotdot* framework for image acquisition and data analysis

We first introduce the *dotdotdot* framework, which involves 1) image acquisition using confocal microscopy or spectral imaging/linear unmixing, 2) image processing to extract nuclei/regions of interest (ROIs) and quantitative transcript abundances and 3) transcript colocalization analysis to classify cell types and 4) differential expression analysis. To localize and quantify single transcripts in individual nuclei, we developed parallel gene-labeling, fluorescence microscopy, and image analysis workflows for mouse and human brain tissues. The general workflows for mouse **(Fig. 1a)** and human **(Fig. 1b)** tissues are similar, but they include optimized conditions for sample preparation (i.e. section thickness, fixation, protease treatment), smFISH labeling (i.e. V1 vs. V2 RNAscope Multiplex Fluorescence Technology, number of gene targets, fluorophores), fluorescent imaging (i.e. confocal microscopy vs. multispectral imaging/linear unmixing), and image analysis (segmentation, dot/transcript detection). Differences in image processing and data analysis workflows in mouse (**Fig. S1**) and human (**Fig. S2**) tissues arose from the need to address a challenge specifically associated with fluorescent imaging in postmortem human brain tissue-lipofuscin autofluorescence. In addition, because postmortem human brain is a limited resource, we sought to maximize multiplexing capabilities by utilizing V2 4-plex RNAscope technology, which allowed us to visualize an additional gene target compared to the V1 3-plex technology used for mouse tissues. For the human workflow (**Fig. 1b**), we used four different fluorophores (Opal520, Opal570, Opal620, and Opal690) to label four distinct gene targets. Importantly, the number associated with each Opal dye corresponds to its maximum emission wavelength ([520nm] green, [570nm] orange, [620nm] red, [690nm] far red, respectively). Following image acquisition, raw fluorescent data are processed in MATLAB using *dotdotdot*. This toolbox and example vignettes are available at: https://github.com/LieberInstitute/dotdotdot. We demonstrate the utility of this framework (**Fig. 2**) and software using several experimental examples across diverse applications in mouse and human tissues.

**Figure 1.**
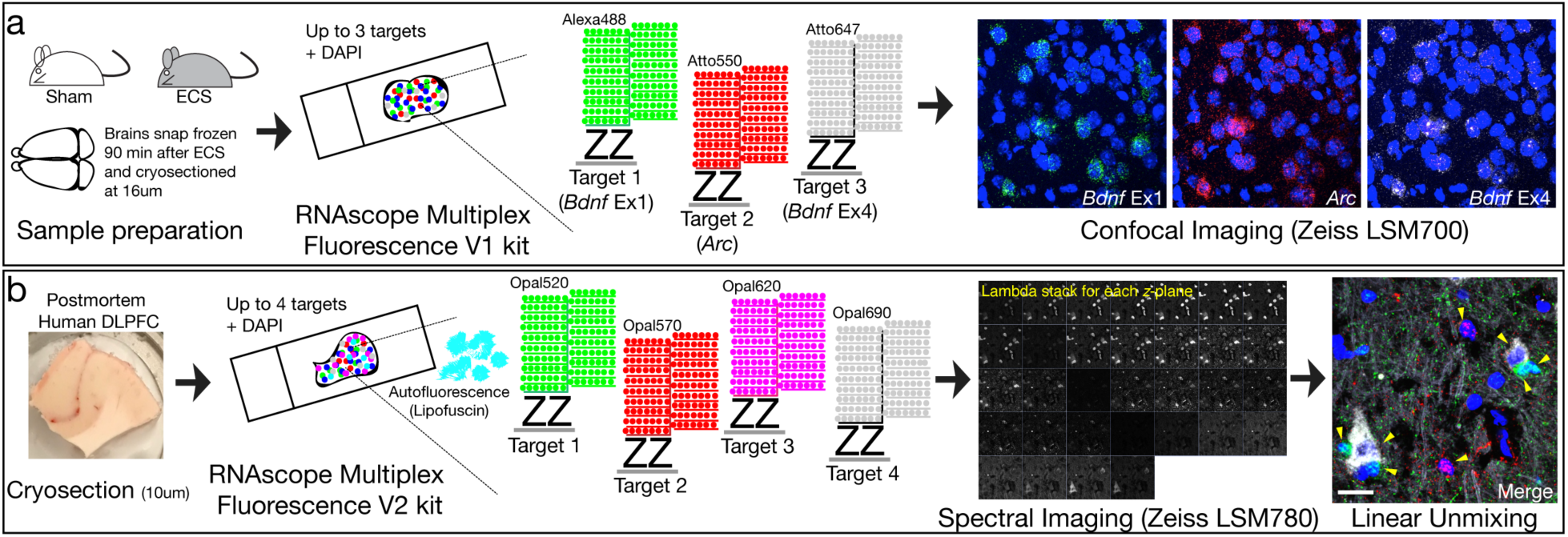
Experimental workflows and imaging protocols for smFISH in mouse and human tissues. **a**, Brains were extracted from wild-type (WT) mice 90 minutes following electroconvulsive seizures (ECS) or Sham treatment and sectioned on a cryostat. Gene targets were visualized with the RNAscope Multiplex Fluorescence V1 kit. RNAscope technology uses hybridization of two independent probes (double Z probes), referred to as a “ZZ pair,” that must bind to the target sequence in tandem for signal amplification to proceed via the subsequent binding of preamplifiers, amplifiers, and fluorescent detection molecules. Approximately 5-30 ZZ pairs are designed for each target gene. After completion of the RNAscope V1 assay, slides are imaged in x, y, and z-dimensions using confocal microscopy. **b**, Fresh frozen postmortem human tissue was sectioned on a cryostat and gene targets were visualized using the RNAscope Multiplex Fluorescence V2 kit. The V2 assay uses the same RNAscope technology with added TSA technology for customization of dyes/concentrations and the ability to include a fourth gene target. The V2 assay is also better suited for tissues with autofluorescence, such as postmortem human brain tissue, which contains an abundance of highly autofluorescent lipofuscin granules. Multispectral imaging and linear unmixing were used to separate individual probe signals and lipofuscin autofluorescence. Lipofuscin signals served as a mask during downstream analysis to exclude pixels confounded by autofluorescence.

**Figure 2.**
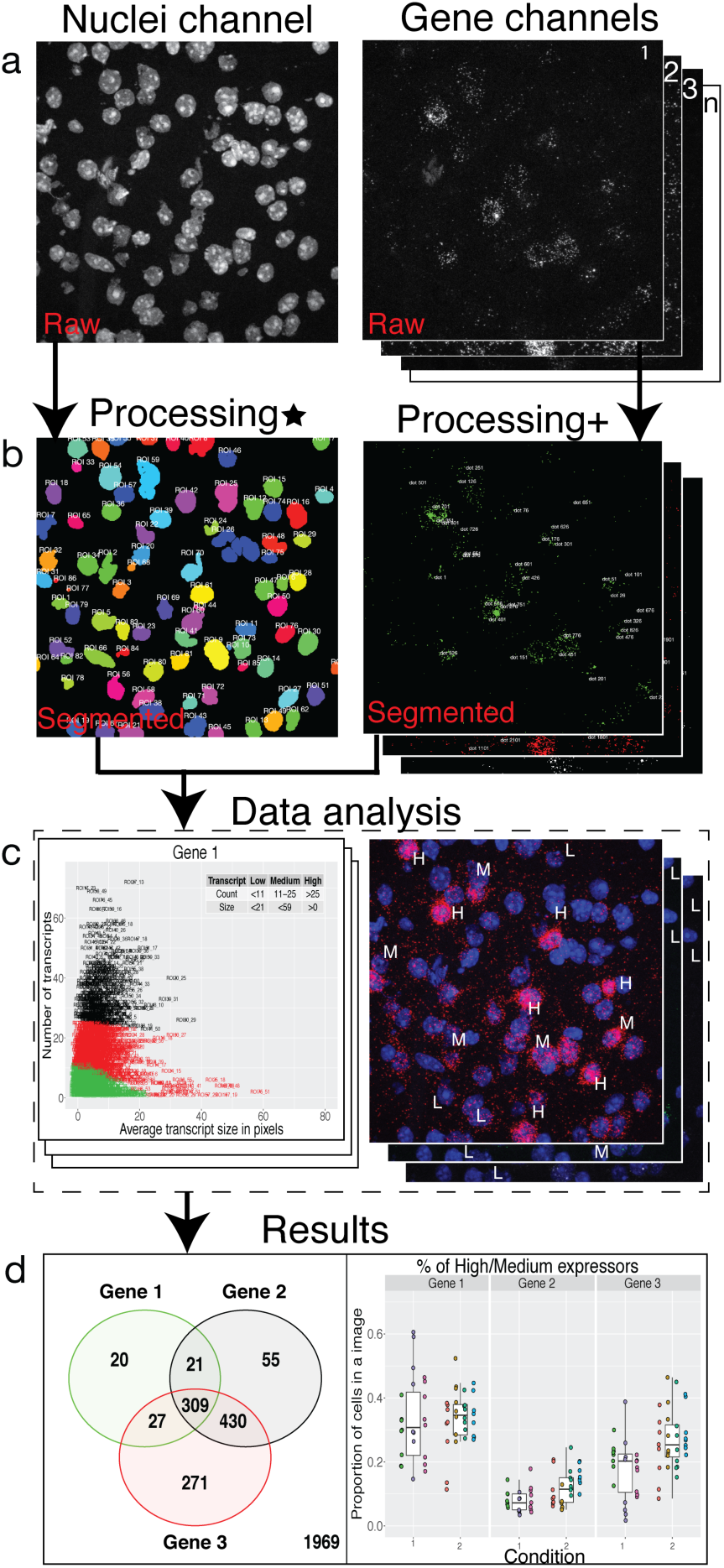
*Dotdotdot* image processing and data analysis workflow: **a**, Raw *‘.czi’* images of nuclei and gene channels. **b**, Final segmented images of nuclei and gene channels (Processing* involves gaussian smoothening, adaptive thresholding and watershed segmentation of nuclei channel, Processing+ involves background filtering, histogram-based thresholding and watershed segmentation of transcript channel). **c**, Data analysis steps (for example, k-means) that are executed based on metrics from segmented images. **d**, Predictions from data analysis steps are used to produce final results.

### *dotdotdot* quantifies dynamics of two activity-regulated genes (ARGs) at cellular resolution following induction of widespread neural activity

We first conducted a proof-of-concept experiment in mouse tissue examining the expression of two well-established activity-regulated genes (ARGs) in piriform cortex following brain stimulation to establish and validate the robustness of *dotdotdot* for quantifying smFISH data acquired with RNAscope technology. Mice were administered electroconvulsive seizures (ECS) to induce widespread neural activity and subsequent upregulation of ARGs^29^. After 90 minutes, brains from ECS- and Sham-treated animals were collected, snap frozen, and processed through our smFISH workflow for mouse tissue to visualize individual transcripts for activity regulated cytoskeleton associated protein (*Arc*) and brain-derived neurotrophic factor (*Bdnf*) splice variants (**Fig. 1a**). *Bdnf* transcription is initiated from one of nine promoters upstream of individual 5′-untranslated regions (UTRs) that are spliced to a common coding exon^30, 31^. We previously designed and validated RNAscope probes for *Bdnf* splice variants containing untranslated exons 1 (Ex1) and 4 (Ex4), which are strongly induced by neural activity^32–34^. As expected, confocal images show a qualitative upregulation of *Arc*, *Bdnf* Ex1, and *Bdnf* Ex4 transcripts following induction of neural activity (**Fig. 3a-b**).

**Figure 3.**
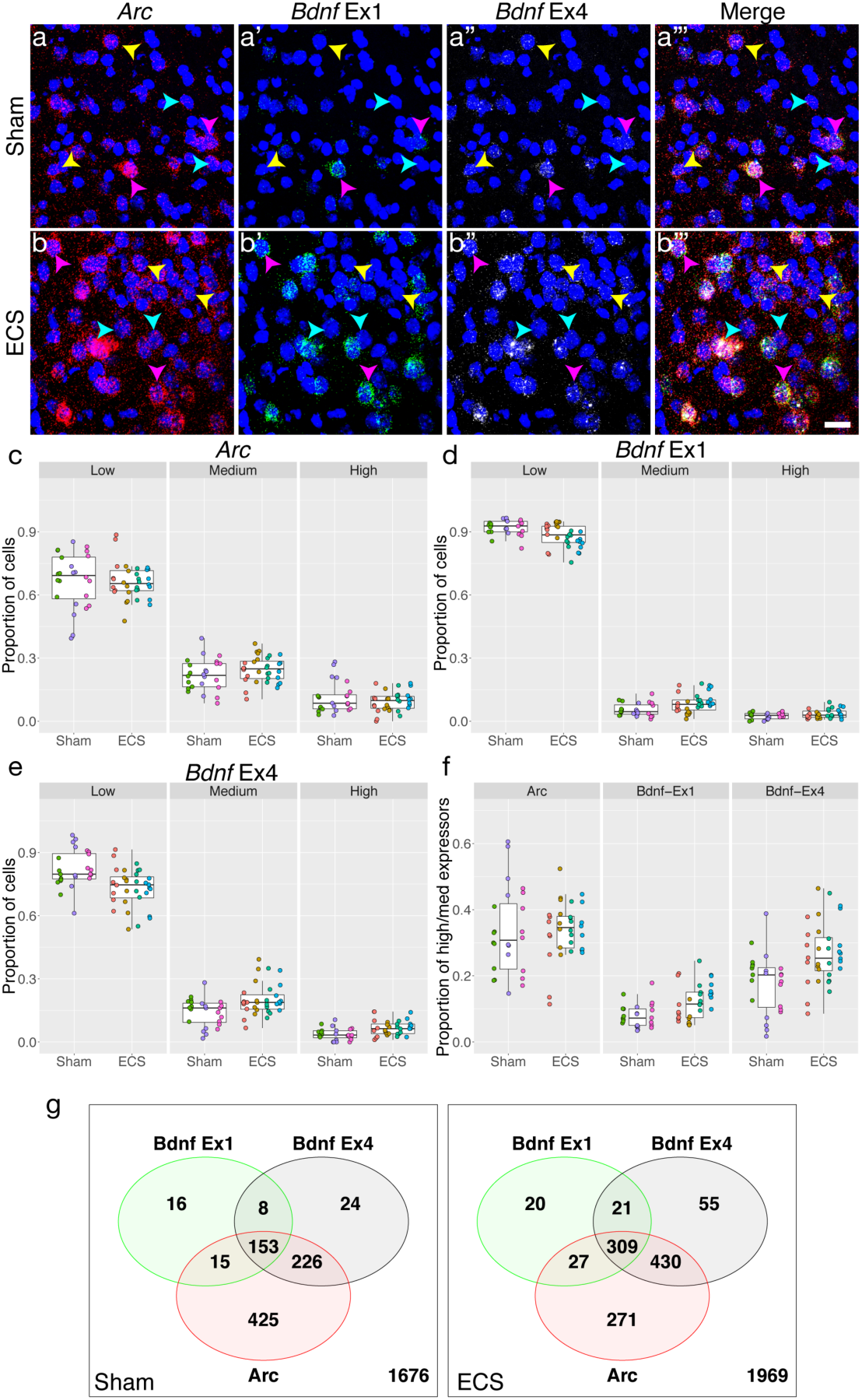
Image analysis with *dotdotdot* captures differential expression of two activity-regulated gene (ARG) transcripts in single cells of mouse cortex following activity induction. **a**, Maximum intensity confocal projections of piriform cortex depicting expression of transcripts for *Arc* (**a**), *Bdnf* Ex1, (**a’**) *Bdnf* Ex4 (**a’’**), and merged (**a’’’**) from a mouse receiving Sham treatment. **b**, Maximum intensity confocal projections of piriform cortex depicting expression of transcripts for *Arc* (**b**), *Bdnf* Ex1, (**b’**) *Bdnf* Ex4 (**b’’**), and merged (**b’’’**) 90 minutes after an acute ECS treatment. **c-e**, Proportion of cortical cells expressing low, medium and high levels of *Arc* (**c**), *Bdnf* Ex1 (**d**), and *Bdnf* Ex4 (**e**) in Sham vs. ECS treatment (n= 3 mice, 24 images, 2543 nuclei and n=4 mice, 32 images, and 3102nuclei, respectively). **f**, Proportion of cortical cells expressing medium/high levels of *Arc*, *Bdnf* Ex1, *Bdnf* Ex4 following ECS treatment. **g**, Venn diagrams showing co-expression of different ARGs in high/medium expressers following Sham or ECS (n=867 high/medium ROIs out of 2543 total ROIs with n=1676 low expressers excluded for Sham, n=1133 high/medium ROIs out of 3102 total ROIs for ECS with n=1969 low expressers). Yellow arrows highlight cells preferentially expressing *Bdnf* Ex4 compared to *Bdnf* Ex1. Cyan arrows highlight cells expressing *Arc*, but not *Bdnf*. Pink arrows highlight cells enriched in *Bdnf* Ex1, which often co-express *Bdnf* Ex4 and *Arc*. Scale bar is 20um.

To quantitatively analyze increases in these ARG transcripts, *dotdotdot* first uses nuclear segmentation in *x*, *y*, and *z*-dimensions to define nuclei regions of interest (ROIs) based on DAPI staining (**Fig. S1, S3a-c)**. As expected, quantification of nuclei/ROI number and size reveals similar metrics between Sham and ECS images (nuclei size: p=0.7, total nuclei: p=0.47; **Fig. S3d-e)**, demonstrating accurate and effective automated three-dimensional nuclear segmentation. After defining ROIs, *dotdotdot* next performs transcript segmentation for each gene in *x*, *y*, and *z*-dimensions (**Fig. S1, S4a-l**). Metrics such as dot location, size, number, and fluorescence intensity are extracted for each transcript. Using dot count metrics, analysis of total, nuclear, and cytoplasmic *Arc*, *Bdnf* Ex1, and *Bdnf* Ex4 transcripts per image reveals increases in these ARGs following ECS (**Fig. S4m-o**). Quantification revealed significant increases in *Bdnf* Ex1 and Ex4 transcripts in both nuclear and non-nuclear (cytoplasmic) compartments (*Bdnf* Ex1 nuc: p=5.05e-4, cyt: p=9.53e-4; *Bdnf* Ex4 nuc: p=1.63e-4, cyt: p=0.022) following ECS administration. Interestingly, we see specific increases in cytoplasmic *Arc* transcripts following activity induction, which is consistent with the role of *Arc* as a cytoskeletal protein (nuc: p=0.782, cyt: p=0.0658)^35^.

While increases in these ARGs following induction of widespread neural activity have been appreciated for decades, two questions have remained outstanding. First, are global increases in activity-induced gene transcription mediated by small increases from many cells or large increases from a select group of “expressers?” Second, do individual neurons differentially express and utilize distinct *Bdnf* splice variants as their source of activity-dependent BDNF? Definitively answering these questions requires quantifying transcript levels at single cell resolution. Using metrics for dot count and size, we performed k-means cluster analysis and classified cells into low, medium, and high expressers for each ARG (**Fig. S5**). For example, *Bdnf* Ex1 low expressers had <11 dots (transcripts) with an average dot size of less than 17 pixels, while *Bdnf* Ex1 high expressers had >25 dots. We then examined the proportion of low, medium, and high expressers between Sham and ECS treatment (**Fig. 3c-e**). For both ARGs, we saw shifts from the proportions of low to medium and high expressers following activity-induction. This was especially true for *Bdnf* Ex 4 (low: p=0.00717, medium p=0.0135, high p=0.00984, **Fig. 3e**). Pooling the proportion of high and medium expressers further demonstrated increased *Bdnf* transcription in single cells following ECS (**Fig. 3f,** *Bdnf* Ex1: p=0.0122, *Bdnf* Ex4 p=0.00717). Surprisingly, co-expression analysis of these ARGs in high and medium expressers following Sham or ECS showed that *Arc* and *Bdnf* splice variants are differentially expressed in single cells at baseline and following activity (**Fig. 3g**). While there is extensive overlap among ROIs expressing *Arc*, *Bdnf* Ex1, and *Bdnf* Ex4, there are several ROIs that express only one or both transcripts suggesting that these ARGs can be dynamically regulated in single cells. These data validate the robustness of *dotdotdot* for transcript segmentation and quantification, and illustrate its utility to provide novel biological insights by analyzing data at cellular resolution.

### Visualization and quantification of single transcripts in postmortem human brain tissue using spectral imaging, linear unmixing, and *dotdotdot*

Extensive efforts are underway to more fully characterize the human brain transcriptome within and across cell types to better understand changes in RNA expression associated with brain development and aging, developmental or psychiatric brain disorders, and local genetic variation. Many of these studies incorporate smFISH in postmortem human brain to validate RNA-seq findings. However, postmortem human brain tissue contains abundant lipofuscin, a highly autofluorescent product of lysosomal digestion that confounds quantification of smFISH signals^13, 14, 23^. To address this problem, we employed multispectral imaging and linear unmixing to isolate and exclude lipofuscin autofluorescence from analysis (**Fig. 4a-c**). In addition to allowing for isolation of lipofuscin autofluorescence, this strategy also allows precise separation of spectrally overlapping fluorophores (i.e. orange [Opal570] and red [Opal620]), which is necessary for utilizing 4-plex technology.

**Figure 4.**
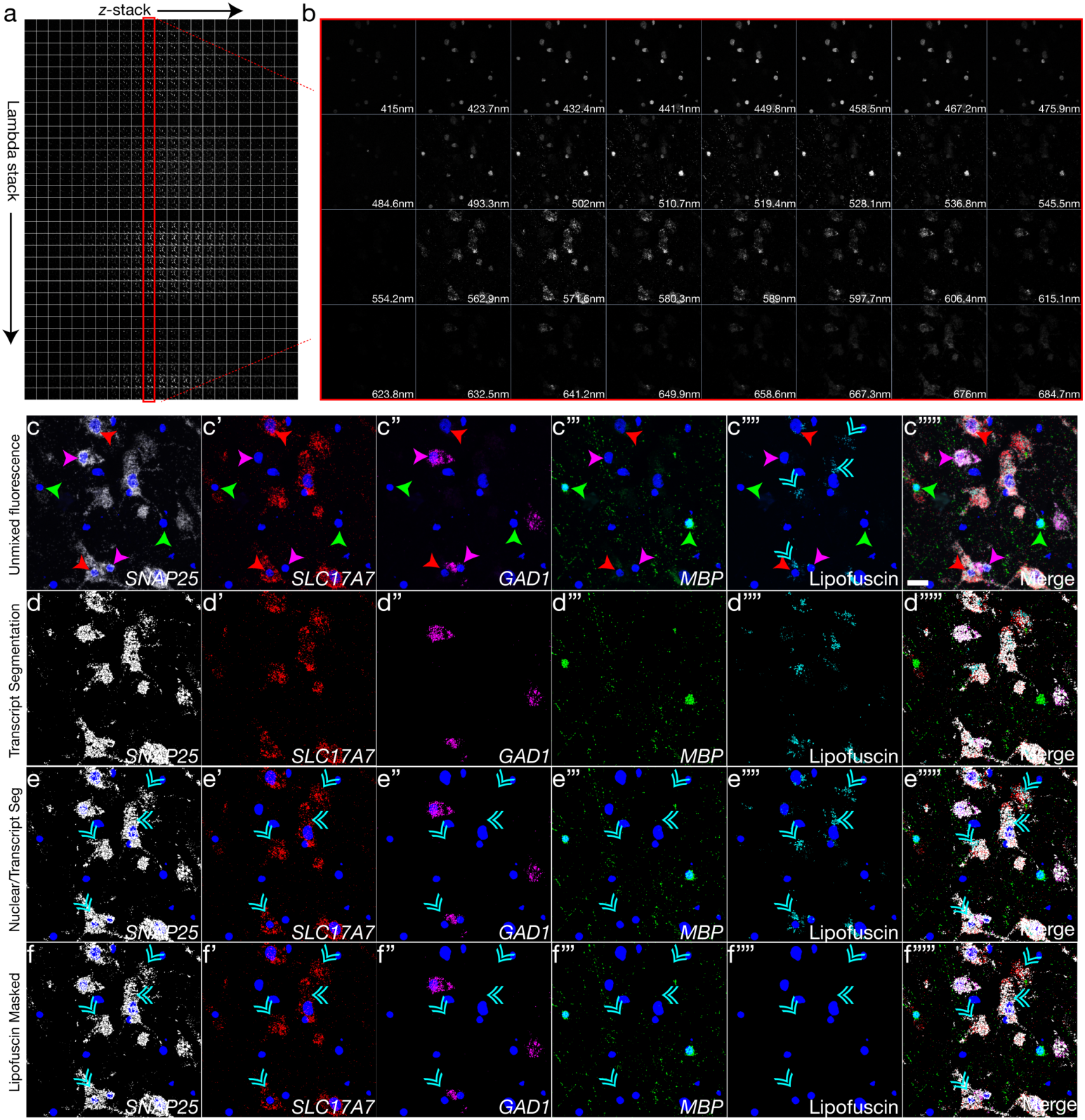
Visualization and quantification of single transcripts in postmortem human brain tissue using spectral imaging/linear unmixing and *dotdotdot*. **a**, Matrix of raw confocal images acquired during spectral imaging in *z*-series of a single field of postmortem human cortex at 63x magnification. A lambda stack is captured at each z-plane detecting expression of *SNAP25*, *SLC17A7*, *GAD1*, *MBP* (labeled using Opal520 [emission maximum at 520nm], Opal570 [emission maximum at 570nm], Opal620 [emission maximum at 620nm], and Opal 690 [emission maximum at 5690nm] dyes, respectively), and lipofuscin autofluorescence. **b**, Representative lambda stack depicting a single z-plane acquired at different wavelength bands, each spanning a limited spectral region (∼8.7nm). **c**, Combined emission signals across the lambda stack in each z-plane are linearly unmixed using reference emission spectral profiles from each Opal dye and lipofuscin to separate the contribution of individual fluorescent gene probes. Unmixed data is then projected across the z-axis. Single transcripts for *SNAP25* (c), *SLC17A7* (**c**’), *GAD1* (**c**’’), and *MBP* (**c**’’’) (canonical markers for neurons, excitatory neurons, inhibitory neurons, and oligodendrocytes, respectively) can be separated from each other and from lipofuscin autofluorescence (**c**’’’’). **d**, Segmentation of unmixed fluorescent signals using *dotdotdot*. **e**, Nuclear segmentation overlaid with transcript segmentation. **f**, Masking with lipofuscin signal removes pixels confounded by autofluorescence from analysis. Green arrows highlight *MBP*+/*SNAP25*-/*SLC17A7*-/*GAD1*- oligodendrocytes. Red arrows highlight *MBP*-/*SNAP25*+/*SLC17A7*+/*GAD1*- excitatory neurons. Pink arrows highlight *MBP*-/*SNAP25*+/*SLC17A7*-/*GAD1*+ inhibitory neurons. Cyan double arrows highlight lipofuscin. Scale bar is 20um.

To validate spectral imaging and linear umixing parameters in postmortem human brain, we used probes targeting canonical cell type markers in dorsolateral prefrontal cortex (DLPFC), including synaptosome associated protein 5 (*SNAP25)*, solute carrier family 17 member 7 (*SLC17A7)*, glutamate decarboxylase 1 (*GAD1)*, and myelin basic protein (*MBP)*, which identify neurons, excitatory neurons, inhibitory neurons, and oligodendrocytes, respectively^3^ (**Fig. 4**). For fluorescent visualization, we assigned Opal690 to *SNAP25*, Opal570 to *SLC17A7*, Opal620 to *GAD1*, and Opal520 to *MBP* and we co-labeled samples with DAPI (maximum emission wavelength at 461nm). We then performed spectral imaging across several *z* planes to generate a matrix of mixed fluorescent signals across the tissue depth (**Fig. 4a**). For a given z-plane, we captured a spectral image stack, or a lambda stack, which is a collection of images of the same field of view (*x*, *y*) captured at different wavelengths (**Fig. 4b**). This four-dimensional matrix of mixed fluorescent signals (*x*, *y*, *z*, lambda stack; **Fig. 4a**) was decoded, or “unmixed,” (**Fig. 4c**) using a linear unmixing algorithm in Zen software, which separates signals from individual probes and lipofuscin autofluorescence using reference emission spectral profiles, or emission “fingerprints,” for each fluorophore (**Fig. S6**) and lipofuscin (**Fig. S7**). A single Opal dye, regardless of its degree of spectral overlap with other Opal dyes and DAPI, has a unique spectral signature that can be cataloged and used to assign the spatial contribution of that fluorophore to individual pixels in a lambda stack during linear unmixing.

Given that reference emission spectral profiles are critical for accurate unmixing, we carefully generated and validated fingerprints for DAPI (**Fig. S6a**), Opal520 (**Fig. S6b**), Opal570, (**Fig. S6c**), Opal620 (**Fig. S6d**), and Opal690 (**Fig. S6e**) in mouse tissue, which lacks lipofuscin autofluorescence. Fingerprints were created in Zen software for each of the Opal fluorophores using a series of 4 “single positive” slides of mouse brain tissue hybridized with a positive control probe against the “house-keeping” gene, *POLR2A*. For the DAPI fingerprint, mouse brain tissue was subjected to pretreatment conditions, but no additional probe labeling before incubation with DAPI. Linear unmixing of single positive slides with all fingerprints (DAPI, Opal520, Opal570, Opal620, and Opal690) verified that reference emission spectral profiles are highly specific for the targeted fluorophore. For example, when *POLR2A* is labeled with the Opal570 fluorophore (**Fig. S6c**), unmixing with the Opal570 fingerprint captures Opal570 fluorescence, while no fluorescent signals are captured with other fingerprints, including the spectrally overlapping Opal620 fingerprint. Transcript segmentation with *dotdotdot* similarly captures fluorescent signals in the appropriate spectral range for each single positive slide. Quantification of dot count, intensity, and size further demonstrates the specificity of reference emission spectral profiles used for linear unmixing of lambda stacks (**Fig. S6**).

In addition to validating robust emission fingerprints for DAPI and each Opal dye, we also generated and validated a spectral signature for lipofuscin autofluorescence in postmortem human DLPFC (**Fig. S7a-b**). Here, we hybridized DLPFC tissue from a representative subject with a negative control probe against the bacterial gene *dapB*. As there was no probe binding, fluorescent signals were attributed exclusively to lipofuscin autofluorescence, and a lipofuscin fingerprint was created in Zen software. Using fingerprints for DAPI, Opal520, Opal570, Opal690, and lipofuscin, we performed linear unmixing of 8 lambda stacks acquired from negative control slides from 4 different subjects (**Fig. S7c**). The lipofuscin fingerprint was equally effective in detecting lipofuscin autofluorescence across subjects. For all images, segregated lipofuscin signals were used to successfully mask and excludes pixels confounded by autofluorescence across the electromagnetic spectrum (**Fig. S7c**).

After linear unmixing with our validated reference emission spectra, we qualitatively observed co-localization of *SLC17A7* (excitatory neurons) and *GAD1* (inhibitory neurons) with *SNAP25* (pan-neuronal) as expected. Similarly, we saw no overlap of *MBP* (oligodendrocytes) with *SNAP25* (neurons), and exclusive expression of either *SLC17A7* (excitatory) or *GAD1* (inhibitory) in *SNAP25*+ neurons (**Fig. 4c)**. Nuclear and transcript segmentation with *dodotdot* faithfully represented fluorescence signals (**Fig. 4d-e**), and masking with lipofuscin signals removed background autofluorescence (**Fig. 4f**). For quantitative analysis of unmixed spectral data using *dotdotdot*, we acquired a set of images at 63x magnification in layers II/III and VI of postmortem DLPFC (n=2 subjects, n=2 cortical strips per subject, n=6 images per layer per strip, **Fig. 4c, 5a-g**). Given differences in the density and arrangement of nuclei in mouse and human tissue, we first optimized nuclear segmentation for human tissue (**Fig. S8a-c**), and demonstrated successful identification of nuclei/ROIs. We observed a similar number and size of nuclei in layer II/III versus layer VI of DLPFC (**Fig. S8d-e,** nuclei number p = 0.376, nuclei size p = 0.592).

Following transcript segmentation and lipofuscin masking with *dotdotdot*, we quantified the number, size, and fluorescence intensity of *SNAP25*, *MBP*, *SLC17A7* and *GAD1* dots and used these data to predict the cell type of each nucleus/ROI using a classification and regression tree (CART) model (**Fig. S9**). Given the close spatial proximity of nuclei/ROIs that may represent different cell types (i.e. *MBP*+ oligodendrocytes situated next to *SLC17A7*+ neurons), this type of model accommodates potentially conflicting signals (i.e. cytoplasm of *MBP*+ oligodendrocytes overlapping the nuclear ROI of a *SLC17A7*+ neuron) allowing accurate cell type calling. We gave the model 60 random nuclei/ROIs from 11 manually annotated images as training data and built a classification tree for defining *SLC17A7*+, *MBP*+, *GAD1*+, or triple negative ROIs (**Fig. S9a**). The confusion matrix for all 201 manually annotated ROIs demonstrates high accuracy between predicted and actual (manual) cell type calling (182 correctly predicted out of 201; **Fig. S9b**). For example, the classifier identifies 31 *MBP*+ ROIs out of 35 actual *MBP*+ ROIs. Of these 31 identified MBP+ ROIs, 30 are correctly predicted and 1 is mislabeled as negative. Plotting the manual and predicted cell type for each ROI against the proportion of the ROI positive for *GAD1*, *SLC17A7*, and *MBP* demonstrates the high sensitivity, specificity, and precision of the classifier, which has a 90.5% accuracy (**Fig. S9c**). Furthermore, for the 961 ROIs identified in layer II/III and layer VI, we confirmed that cells predicted to be excitatory and inhibitory neurons (*SLC17A7*+ and *GAD1*+, respectively) were highly enriched for the pan-neuronal marker *SNAP25* (*SNAP25* vs. Neg, p = <2e-16; *SNAP25* vs. MBP, p = <2e-16 **Fig. 5h**). Consistent with a predicted glial cell type, ROIs classified as oligodendrocytes (MBP+) or negative for *MBP*, S*LC17A7*, and *GAD1* (likely astrocyte and microglia populations) showed low or zero levels of *SNAP25*. Further supporting the accuracy of transcript quantification with *dotdotdot* and subsequent cell type calling with CART analysis, we observed a higher proportion of excitatory neurons compared to inhibitory neurons in both layers of DLPFC and a higher proportion of oligodendrocytes in layer VI compared to layer II/III (**Fig. 5i,** *MBP:* p = 3.97e-05, *SLC17A7:* p = 0.029, *GAD1:* p = 0.0493, Neg: p = 0.859874). Taken together, we show robust analysis of gene expression in postmortem human DLPFC using 4-plex RNAscope technology and spectral imaging/linear unmixing in combination with *dotdotdot* and machine learning.

**Figure 5.**
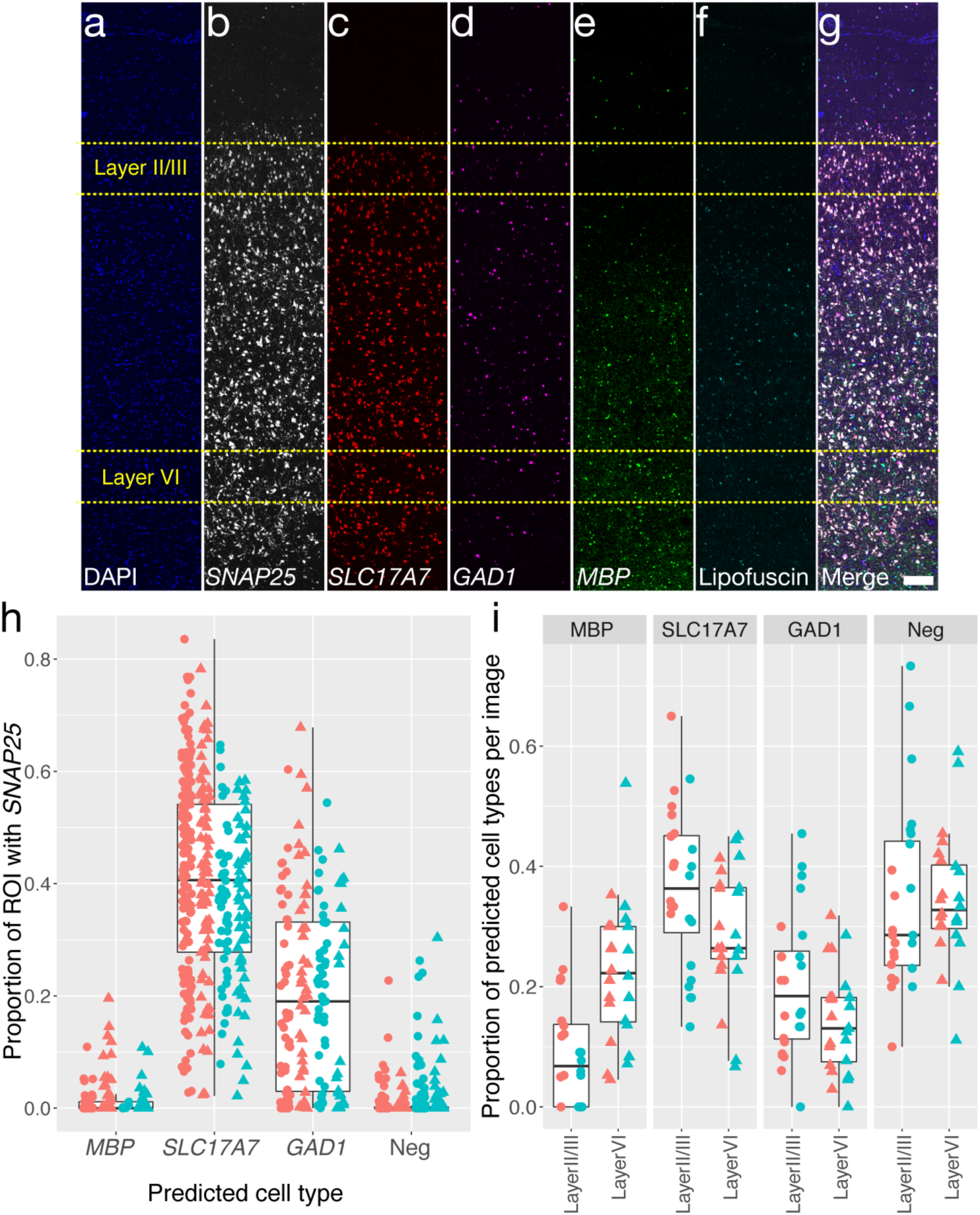
Layer-specific cell type analysis in postmortem human DLPFC demonstrates sensitivity and specificity of *dotdotdot* for transcript quantification. **a-g**, Maximum *z*-projection of unmixed and stitched lambda stacks acquired across a single cortical strip (layer I to VI) of postmortem human DLPFC depicting expression of DAPI (**a**), *SNAP25* (**b**), *SLC17A7* (**c**), *GAD1* (**d**), *MBP* (**e**), lipofuscin (**f**) and merged (**g**). High magnification (63x) images used for analysis were randomly acquired within a strip in layer II/III (80-90% of strip thickness; n= 24 images total) or layer VI (20-30% of strip thickness; n=24 images total) from 2 individuals from 2 different strips (see Fig. 3 for representative 63x image). **h**, Validation of cell type calling using CART analysis demonstrating an enrichment of *SNAP25* transcripts in predicted excitatory (*SLC17A7*) and inhibitory (*GAD1*) neurons compared to oligodendrocytes (*MBP*) and *SLC17A7*-/*GAD1*-/*MBP*-cells (likely astrocytes and microglia). Color represents different subjects and shape represents different layers (circle=layer II/III and triangle is layer VI). **i**, Proportion of cortical cells expressing markers for neurons, excitatory neurons, inhibitory neurons, and oligodendrocytes in layer II/III vs. layer VI of DLPFC (n= 519 ROIs layer II/III and n= 442 ROIs layer VI). Scale bar is 200um.

### *dotdotdot* accommodates smFISH data acquired with alternative imaging acquisition parameters and saved in diverse file formats

To demonstrate the utility of our image acquisition and analysis workflow, we enabled flexible solutions for users who may be unable to image lipofuscin autofluorescence with spectral imaging or who may acquire images in alternative file formats. First, we evaluated whether a narrower spectral range, such as the range detected by the Opal520 fingerprint, could be used for lipofuscin detection and masking. This would allow users to sacrifice data collection for one gene target in order to capture lipofuscin autofluorescence in the available spectral range (i.e. use the spectral range of one fluorophore to capture lipofuscin instead of a gene target).

To evaluate this possibility, we hybridized postmortem human DLPFC with probes targeting *SLC17A7* (Opal690), *GAD1* (Opal620), and *SNX19* (sorting nexin 19, Opal570), a gene associated with genetic risk for schizophrenia^36^ (**Fig. S10**). As the Opal520 fluorophore was not used to detect a probe target, signals unmixed with the Opal520 fingerprint represented autofluorescence due to lipofuscin (**Fig. S10a**). Unmixing with the Opal520 fingerprint as a proxy for the lipofuscin fingerprint captured the majority of autofluorescence. However, unmixing the same lambda stack with the custom lipofuscin fingerprint was more comprehensive and captured a larger number of autofluorescent pixels (**Fig. S10b**). Quantification of fluorescent pixels assigned to each gene demonstrated that more confounding autofluorescent pixels were removed from gene channels when unmixing with the lipofuscin compared to Opal520 fingerprint (**Fig. S10c**). However, masking with either Opal520 or lipofuscin unmixed signals removed a substantial portion of background autofluorescence (**Fig. S10a-b**), suggesting that a more narrow spectral range can be used to detect and monitor lipofuscin autofluorescence if imaging capabilities are limited.

Finally, given the diversity of imaging file formats for smFISH data, we next evaluated the performance of *dotdotdot* on images acquired and saved using a different microscope system and customized file format (Nikon “*.nd2”* files vs. Zeiss “.*czi”* files). For three distinct combinations of probe targets hybridized to postmortem human cortical tissue (images courtesy of Jennie Close and Ed Lein at the Allen Brain Institute), we show accurate segmentation of fluorescent RNAscope signals (**Fig. S11**). *dotdotdot* is compatible with several additional file formats, including those supported by Zeiss, Leica, Nikon, Olympus, and MetaMorph systems. Taken together, we provide a robust, flexible, and automated approach for quantification of multiplex single molecule fluorescent in situ hybridization (smFISH) images in complex tissues, including postmortem human brain.

## DISCUSSION

Here we present and validate *dotdotdot,* a versatile computational tool for quantitative analysis of smFISH data. We demonstrate robustness of this framework for quantifying gene expression in single cells in mouse brain as well as postmortem human brain. Importantly, we develop imaging acquisition and analysis strategies for detecting and removing tissue autofluorescence in smFISH data to improve the accuracy of transcript quantification. Furthermore, we demonstrate that quantitative segmentation data can be used in combination with machine learning approaches to define cell types in a systematic and unbiased manner. Finally, we demonstrate the flexibility of *dotdotdot* for diverse data file formats and imaging acquisition parameters thereby increasing its utility for different types of smFISH data.

### *dotdotdot* confers advantages for quantitative analysis of smFISH data

*dotdotdot* is compatible with several smFISH workflows, including RNAscope Multiplex Fluorescent V1 and V2 assays (**Fig. 1**). We demonstrate analysis of six separate fluorescent signals using spectral imaging and linear unmixing (**Fig. 5**), but there is no limit to the number of gene targets or fluorescent signals (i.e. DAPI or lipofuscin) that *dotdotdot* can process in parallel. Chemistry for higher order multiplexing is rapidly coming online. For example, Advanced Cell Diagnostics recently introduced the RNAscope Hi-plex assay, which visualizes up to 12 gene targets in a single tissue slice with several rounds of hybridization and stripping. Computational tools like *dotdotdot* will be urgently needed to quantitatively analyze multiple fluorescent signals with a rapid and unbiased approach.

While several commercial platforms, including HALO (Indica Labs) and Aperio (Leica), are available for analysis of smFISH data, these softwares are only available for purchase and they utilize proprietary algorithms that offer less flexibility for customization. *dotdotdot* is an open source platform that can be implemented by any MATLAB user and can accommodate diverse smFISH data and file formats. Furthermore, unlike other open-source platforms^12^, *dotdotdot* is optimized for complex tissues, and can process three-dimensional data (i.e. *z*-stacks) by segmenting each *z*-plane as opposed to two-dimensional maximum intensity projections. This strategy allows for accurate quantification of fluorescent signals across tissue depths in complex biological samples.

### Quantifying spatial gene expression at single cell resolution with *dotdotdot* delivers novel biological insights

We confirmed the accuracy of *dotdotdot* by quantifying expression of *Arc* and *Bdnf,* ARGs that are expected to increase following ECS administration^29, 37–40^. While induction of *Arc* and *Bdnf* in response to ECS has been long appreciated^41^, it has remained unclear whether activity-induced increases are driven by increased transcription in a stable population of cell “expressers,” recruitment of additional neurons that become transcriptionally active, or a combination of both. Furthermore, whether multiple *Bdnf* isoforms can be induced within an individual BDNF-expressing cell *in vivo* has been difficult to demonstrate. While the necessary spatial and cellular resolution to answer these questions is lost in bulk homogenate analysis of gene expression, it can be resolved with quantitative analysis of smFISH data.

Using *dotdotdot*, we identify distinct ensembles of ARG-expressing cells following ECS. While the majority of cells upregulate *Arc* and *Bdnf* upon activity, we identify some cells that appear to selectively upregulate *Arc*, and some that preferentially upregulate specific *Bdnf* isoforms. The dynamics in regulation of these ARGs following activity have important implications for studies using activity-induced promoters (e.g. *Arc* and *Fos)* to tag ensembles of activated neurons^42, 43^. These data suggest that “activated” ensembles may not be a homogenous population as some activity-tagged cells may have the capacity to release BDNF and engage downstream plasticity cascades, while others may not. Similarly, our results suggest that activity-induced, BDNF-expressing neurons are likely not a uniform population as some BDNF-expressing neurons may preferentially utilize one isoform over another. Because *Bdnf* isoforms show distinct expression kinetics and subcellular targeting^44–46^, it is likely that differences in upregulation between single cells impact neuron function. Future studies should aim to link differences in ARG expression following activity or distinct behaviors to correlates of cell function, including morphology and activity, to better understand how dynamic ARG expression impacts network function.

### *dodotdot* overcomes tissue autofluorescence to accurately profile postmortem human brain tissue

Postmortem human brain poses several intrinsic and extrinsic challenges for quantitative smFISH, including extensive cellular heterogeneity^47^, high lipofuscin autofluorescence^13^, and RNA degradation due to the interval between death and brain extraction and processing. To overcome these challenges, we carefully optimized the RNAscope assay as well as imaging acquisition and analysis parameters. We found that RNAscope Multiplex Fluorescence V2 reagents afforded higher signal-to-noise in postmortem human brain tissue, especially for moderate or low expressing gene targets, such as *SNX19* (**Fig. S10**). Flexibility to dilute fluorophore concentrations for higher expressing gene targets, such as *SLC17A7* (**Fig. 4**), also facilitated more accurate target labeling and quantification. While we utilized RNAscope technology for fluorescent probe labeling, non-commercial smFISH technologies have also proven successful in postmortem human brain tissue^4^. However, RNAscope offers a rapid, robust, and universal approach that has now been replicated by several groups for qualitative validation of RNA-seq data in postmortem human brain tissue^3–5, 18^.

For quantitative analysis of smFISH signals in postmortem human brain tissue, it is necessary to identify and remove fluorescent pixels derived from lipofuscin autofluorescence. There are several lipophilic reagents marketed to quench tissue autofluorescence that can be incorporated into the RNAscope assay^23, 48^. However, dye-based approaches proved ineffective in mitigating lipofuscin signals in our hands. While we did not explore optical tissue clearing methods to remove lipofuscin granules, several approaches, such as CLARITY, have been utilized in conjunction with smFISH^28, 49, 50^ and may be compatible with the RNAScope assay.

Given the persistence of lipofuscin in our human brain samples, we employed a spectral imaging/linear unmixing approach to isolate and mask autofluorescent pixels attributed to lipofuscin and other tissue artifacts^23^. This approach also allowed us to conduct 4-plex labeling as we were able to separate signals arising from spectrally overlapping fluorophores (Opal570 and Opal620). We carefully validated reference emission profiles used for linear unmixing of fluorescent probe signals (**Fig. S6**) and lipofuscin autofluorescence (**Fig. S7**). Importantly, we demonstrated that the spectral quality of lipofuscin autofluorescence is comparable among subjects close in age and a common lipofuscin fingerprint can be employed for multiple subjects. However, studies examining gene expression across development may require age-specific lipofuscin fingerprints as biochemical and autofluorescent properties of lipofuscin are dynamic across the lifespan^14^. Although spectral imaging or a custom lipofuscin filter cube are superior imaging strategies for detecting and masking lipofuscin autofluorescence^21, 23^, we also demonstrate the feasibility of using a narrower spectral range, such as that detected by Opal520 (or a standard FITC filter), to detect and mask lipofuscin fluorescence with *dotdotdot*. Given that neurons are more profoundly affected by lipofuscin masking than glia^13^, it is important to consider whether removal of lipofuscin pixels biases quantification of gene expression in particular cell types. Our data suggest that neurons are not adversely affected by lipofuscin masking as we detect the expected proportions of neuronal and glial cell types in DLPFC^51^ **(Fig. 5**).

### dotdotdot complements growing computational approaches for spatial analysis of genome-wide expression

We developed *dotdotdot* to faithfully segment fluorescent probe signals and provide quantitative information on transcript/dot size, number, and fluorescence intensity. While this quantitative output of *dotdotdot* can stand alone to answer many biological questions^52^, we provide examples of how further computational approaches can be utilized to answer more complex questions about gene expression. First, in mouse tissue, we used K-means clustering ^53^ to identify groups of cells defined as high, medium, and low expressers for individual genes based on transcript dot size and number (**Fig. 3**). Second, using thresholds established with the K-means approach, we examined co-localization of different ARGs to understand co-regulation of activity-induced transcripts. A key advantage of the K-means approach is that several features of transcript segmentation, such as dot size and number, are included to provide a more comprehensive and unbiased interpretation of gene expression. Finally, given the diversity and intermingled spatial position of different cell types in postmortem human brain tissue, we used a machine learning approach (CART),^54^ to systematically assign cell types to individual ROIs with 91% accuracy (**Fig. 5 and S9**). This model accommodated potentially conflicting signals from mutually exclusive canonical cell type markers (i.e. *MBP* and *SLC17A7*) to assign the most likely cell type for each ROI. As molecular profiles of cell types become more complex, machine learning approaches such as CART may become increasingly necessary to interpret overlapping patterns of gene expression^55^.

Spatial analysis of genome-wide expression is a rapidly emerging field^2, 56–58^. With the advent of Spatial Transcriptomics^59^ and Slide-seq^60^, smFISH will continue to be a gold standard for validating spatial RNA-seq approaches. *dotdotdot* is an intuitive computational tool that can add quantitative dimensions to traditionally qualitative smFISH data. Furthermore, as tools for integrating smFISH and single cell RNA-seq data continue to develop^61, 62^, *dotdotdot* can augment these approaches by extracting quantitative information from existing datasets for integration with spatial gene expression databases. In summary, we present a computational tool for smFISH data that can be readily implemented in wet-bench laboratories to elevate spatial analyses of gene expression and complement growing single cell and spatial transcriptomic data sets in the field of neuroscience^63^ and beyond^64^.

## MATERIALS AND METHODS

### Animals and electroconvulsive seizure (ECS) treatment

Six-week old male mice (C57BL/6J) were administered either Sham or ECS treatment as previously described^41, 65^. Briefly, ECS was delivered with an Ugo Basile pulse generator using a corneal electrode fork placed over the frontal cortex (model #57800-001, shock parameters: 100 pulse/s frequency, 0.3 ms pulse width, 1s shock duration and 50 mA current). The stimulation parameters were chosen because they reliably induced tonic-clonic convulsions. Mice were administered inhaled isoflurane anesthesia prior to ECS sessions, and remained anesthetized for the procedure. Each mouse received a single session of Sham or ECS and was euthanized 90 minutes following the treatment. All animal experiments were approved by the SoBran Institutional Animal Care and Use Committee.

### Postmortem human tissue samples

Post-mortem human brain tissue from two donors (both male: one 17 years old of African American ancestry and the other 25 years old with European ancestry) was obtained by autopsy primarily from the Offices of the Chief Medical Examiner of the District of Columbia, and of the Commonwealth of Virginia, Northern District, all with informed consent from the legal next of kin (protocol 90-M-0142 approved by the NIMH/NIH Institutional Review Board). Clinical characterization, diagnoses, and macro- and microscopic neuropathological examinations were performed on all samples using a standardized paradigm, and subjects with evidence of macro- or microscopic neuropathology were excluded. Details of tissue acquisition, handling, processing, dissection, clinical characterization, diagnoses, neuropathological examinations, RNA extraction and quality control measures have been described previously^66^.

### 3-plex smFISH and image acquisition in mouse tissue

Mice (n=3 Sham and n=4 ECS treated) were cervically dislocated and brains were removed from the skull, flash frozen in isopentane, and stored at -80°C. Brain tissue was equilibrated to - 20°C in a cryostat (Leica, Wetzlar, Germany) and serial sections of piriform cortex were collected at 16μm (4 sections per slide). Sections were stored at -80°C until completion of the smFISH assay. For mouse studies (**Fig. 1a**), in situ hybridization assays were performed with RNAscope technology utilizing the RNAscope Fluorescent Multiplex Kit V1 (Cat # 320850 Advanced Cell Diagnostics [ACD], Hayward, California) according to manufacturer’s instructions as previously described^32^. Briefly, tissue sections were fixed with a 10% neutral buffered formalin solution (Cat # HT501128 Sigma-Aldrich, St. Louis, Missouri) for 20 minutes at room temperature (RT), series dehydrated with ethanol, and pretreated with protease IV for 20 minutes. Sections were incubated with custom-designed probes for *Bdnf* exon IV (Cat # 482981-C3, ACD) and commercially probes for *Bdnf* exon I and *Arc* (Cat #457321-C2 and #316911, ACD, Hayward, California). Probes were fluorescently labeled with orange (excitation 550 nm), green (excitation 488 nm), or far red (excitation 647) fluorophores using the Amp 4 Alt B-FL and stained with DAPI (4′,6-diamidino-2-phenylindole) to demarcate the nucleus. Confocal images were acquired in *z*-series using a Zeiss LSM700 confocal microscope. For each mouse (biological replicate), two images were randomly captured in the piriform cortex per section (4 sections; 8 images total).

### 4-plex smFISH and image acquisition in postmortem human tissue

Two blocks of fresh frozen dorsolateral prefrontal cortex (DLPFC) from neurotypical control individuals ages 24 and 17 were sectioned at 10μm and stored at -80°C. RNA integrity numbers (RINS) were 8.4 and 8.8, respectively. For postmortem human studies (**Fig. 1b**), *in situ* hybridization assays were performed with RNAscope technology utilizing the RNAscope Fluorescent Multiplex Kit V2 and 4-plex Ancillary Kit (Cat # 323100, 323120 ACD, Hayward, California) according to manufacturer’s instructions. Briefly, tissue sections were fixed with a 10% neutral buffered formalin solution (Cat # HT501128 Sigma-Aldrich, St. Louis, Missouri) for 30 minutes at RT, series dehydrated in ethanol, pretreated with hydrogen peroxide for 10 minutes at RT, and treated with protease IV for 20 minutes. Sections were incubated with probes for *SNAP25*, *SLC17A7*, *GAD1*, and *MBP* (Cat #518851, 415611-C2, 573061-C3, 573051-C4, ACD, Hayward, California) and stored overnight in 4x SSC (saline-sodium citrate) buffer. Probes were fluorescently labeled with Opal Dyes (Perkin Elmer, Waltham, MA; Opal690 diluted at 1:1000 and assigned to *SNAP25*; Opal570 diluted at 1:1500 and assigned to *SLC17A7*; Opal620 diluted at 1:500 and assigned to *GAD1*; Opal520 diluted at 1:1500 and assigned to *MBP*) and stained with DAPI (4′,6-diamidino-2-phenylindole) to label the nucleus. For experiments with *SNX19*, sections were incubated with probes for *SLC17A7*, *GAD1*, and *SNX19* (Cat #415611-C3, Cat #404031, Cat #518861-C2, ACD, Hayward, California) and stored overnight in 4x SSC buffer. Probes were fluorescently labeled with Opal Dyes (Opal690 diluted at 1:1500 and assigned to *SLC17A7*; Opal570 diluted at 1:500 and assigned to *SNX19*; Opal520 diluted at 1:1000 and assigned to *GAD1*) and stained with DAPI to label the nucleus

Lambda stacks were acquired in z-series using a Zeiss LSM780 confocal microscope equipped with 20x x 1.4 NA and 63x x 1.4NA objectives, a GaAsP spectral detector, and 405, 488, 555, and 647 lasers. All lambda stacks were acquired with the same imaging settings and laser power intensities. For each subject, two cortical strips were tile imaged at 20x to capture layers I to VI (**Fig. 5**). Layer II/II and layer VI were identified by measuring 20-30% and 80-90% of the cortical layer thickness, respectively. This strategy reliability delineated layer II/III and VI across 10 individuals and cortical strips with varying absolute thicknesses. After demarcation of cortical layers, the positions feature in Zen software was used to randomly select 6 fields per layer per strip (n=12 layer II/III and n=12 layer VI in 2 different cortical strips per subject) for high magnification imaging at 63x. Following image acquisition, lambda stacks in *z*-series were linearly unmixed in Zen software (weighted; no autoscale) using reference emission spectral profiles previously created in Zen (see below) and saved as Carl Zeiss Image “*.czi*” files.

### Generation of Reference Emission Spectral profiles

Reference emission spectral profiles, or “fingerprints,” were created for each Opal dye in Zen software. Briefly, 4 single positive slides were generated in mouse tissue using the RNAscope Fluorescent Multiplex Kit V2 and 4-plex Ancillary Kit (Cat # 323100, 323120 ACD, Hayward, California) and a control probe against the housekeeping gene *POLR2A* according to manufacturer’s instructions as described above (**Fig. S6)**. Mouse tissue was used in place of human tissue due to lower tissue autofluorescence (i.e. the absence of confounding lipofuscin signals). For each single positive slide, *POLR2A* was labeled with either Opal520, Opal570, Opal620, or Opal690 dye. A single positive slide was generated for DAPI using the same pretreatment conditions, but omission of probe hybridization steps. To generate a reference emission spectral profile for lipofuscin autofluorescence, a negative control slide was generated in postmortem DLPFC tissue using a 4-plex negative control probe against 4 bacterial genes (Cat #321831, ACD, Hayward, California) in which all Opal dyes were applied, but no probe signal was amplified.

### Automated Imaging Analysis

*dotdotdot* is a MATLAB-based command line toolbox for automated nuclei and transcript segmentation and quantification^67^. Confocal images are processed in MATLAB, but downstream data analyses can be performed in R (as done here), MATLAB, python, or any statistical software. Briefly, the processing pipeline involves smoothing/filtering raw images, thresholding, watershed segmentation, autofluorescence masking, and extracting dot metrics. The analysis pipeline involves k-means clustering for classifying nuclei expression (low, medium and high) and CART (Classification and Regression Trees) for classifying cell types (astrocytes, oligodendrocytes, GABAergic or glutamatergic neurons). Bio-formats toolbox “bfmatlab” ^68^ is used to read the image data into a MATLAB structure with fields containing gene data, DAPI and lipofuscin. Processing techniques for human nuclei, mouse nuclei and transcript channels are different, as described below.

#### Mouse nuclei segmentation

Processing and segmentation of mouse nuclei is performed using the MATLAB toolbox called “CellSegm”^69^. The toolbox provides the user with several input options for smoothing (coherence enhancing diffusion, edge enhancing diffusion, gaussian) and thresholding (iterative thresholding, adaptive thresholding, gradient thresholding, ridge enhancement). A 2D planewise gaussian smoothing with default settings (filter size = [5 5], standard deviation = 2) was used to filter the raw images (X = Y = 512 pixels) (**Fig S1.A1**). Adaptive thresholding with an average filter (size = [42 42]) followed by several morphological operations (like imopen, imerode, imfill) were performed on the gaussian smoothed images (**Fig S1.B1**) to obtain the binary image (**Fig S1.C1**). The irregularly large objects in the binary image are then split into smaller segments using watershed segmentation based on local maxima and the euclidean distances (**Fig S1.D1**).

#### Human nuclei segmentation

A 3D median filter (size = [19 19 3]) is used to smooth (**Fig S2.B1**) the intensity irregularities in the raw image (X = Y = 1024 pixels) (**Fig S2.A1**) that are produced from heterogeneously-stained nuclei. An intensity threshold from the image histogram is then used to segment the DAPI stained nuclei from the background. A technique called “*minima imposition*” is applied to the binary image (**Fig S2.C1**) before watershed transform to filter the tiny local minima that might cause over-splitting of large segmented nuclei blobs. A modified distance transform of the binary image is then computed for the watershed segmentation on the maximum *z* projection (**Fig S2.D1**).

#### Transcript segmentation and lipofuscin masking

Background noise (i.e. potential bleed-through from adjacent wavelengths) in the gene channels is eliminated using the function “*imhmin*” ^70^. Here all the minima in the grayscale image whose depth is less than the standard deviation of the image is suppressed (**Fig S1.B2, Fig S2.B2**). A histogram-based intensity threshold is used to segment the RNA signal (**Fig S1.C2, Fig S2.C2**). Watershed segmentation based on the minima of the image is then performed to split the detected pixel clusters in each channel into identified transcripts (**Fig S1.D2, Fig S2.D2**). Lipofuscin segmentation is similar, except it does not include the background suppression step (**Fig S2.Lipofuscin channel**).

#### Extract nuclei and transcript metrics

Custom MATLAB functions (regionprops3 function in Image Processing toolbox) were then used to calculate relevant metrics (count, size, location, intensity) of detected nuclei and transcripts (**Fig S1.E(1,2), Fig S2.E(1,2)**). For human data, before transcript quantification, the segmented RNA channels (**Fig S2.D2**) are masked (**Fig S2.E3**) with the segmented lipofuscin channel (**Fig S2.C3**). Nuclei and transcript colocalization data (**Fig S1.Data analysis.1, Fig S2.Data analysis.1**) are then obtained by assigning each transcript to a cell based on its position in 3 dimensions. For gene expression analysis (mouse data) a dot is assigned to a nucleus if its center falls within the boundary of the nucleus and for cell type classification (human data) each segmented gene pixel is considered as a transcript and is assigned to nucleus if it is within the boundary of the nucleus.

#### Downstream data analysis

(a) Gene expression analysis (**Fig S1.F**): all the nuclei are clustered into low, medium and high expressers for each gene type by the k-means clustering method (**Fig S1.F.2**) based on the total transcript count and the average transcript size per nuclei. The choice of k here (3) was biologically motivated, but can be a flexible parameter dependent on the underlying study design and research question. Nuclei with at least one transcript are recruited for k-means clustering, and the nuclei with zero transcripts are explicitly labeled as low expressers. (b) Cell type classification (**Fig S2.F**): the proportion of each type of transcript in individual nuclei is used by Classification and Regression Trees (CART) to predict the underlying cell type (**Fig S2.F.3**). The initial CART model was built on test and train dataset created from 89 manually annotated nuclei (60 random ROIs were used to train the model and the rest were used to test) from 5 random images (**Fig S2.F.2**) from the whole dataset. The predictions from this model were used to classify the rest of the data into predefined categories. This strategy can be used to develop other analogous classification models for other cell and tissue types.

### Statistics

All statistical analyses were performed in R^71^. For mouse data analyses, we used linear mixed effects modeling with the lmerTest package^72^ to analyze nuclei size, total number of nuclei in an image, gene expression (*Bdnf* Ex1, *Bdnf* Ex4, Arc) as a function of the treatment main effect (Sham versus ECS) and random intercepts of repeated measures: animal ID, brain section and the image/scan ID. For human data analysis, we also used linear mixed effects modeling for the analysis of nuclei size, total number of nuclei in image, proportions of cell types in an image as a function of brain layer (Layer II/III versus Layer VI) as a main effect and random intercepts of repeated measures: brain donor, strip number, and the image ID. For the validation of *SNAP25* enrichment in *GAD1* and *SLC17A7* positive cells, *GAD1* and S*LC17A7* cells were combined into one group and compared to the *MBP* and negative labelled cells. The analysis was performed using the linear mixed effects model with main effects as the predicted cell labels and random intercepts being the same variables as above.

### Data and Code availability

The source code for *dotdotdot* is available together with test data and detailed tutorials at https://github.com/LieberInstitute/dotdotdot.

## ACKNOWLEDGEMENTS

We are grateful for the contributions of the Clinical Brain Disorders Branch of the National Institute of Mental Health in assisting the Lieber Institute for Brain Development in the acquisition and curation of brain tissue donations for this study. We thank Ed Lein and Jennie Close at the Allen Brain Institute for generously sharing RNAscope image data. We thank Daniel Weinberger and Stephanie Page for comments on the manuscript. Funding for these studies was provided by the Lieber Institute for Brain Development.

## SUPPLEMENTARY FIGURES

**Supplementary Figure 1.**
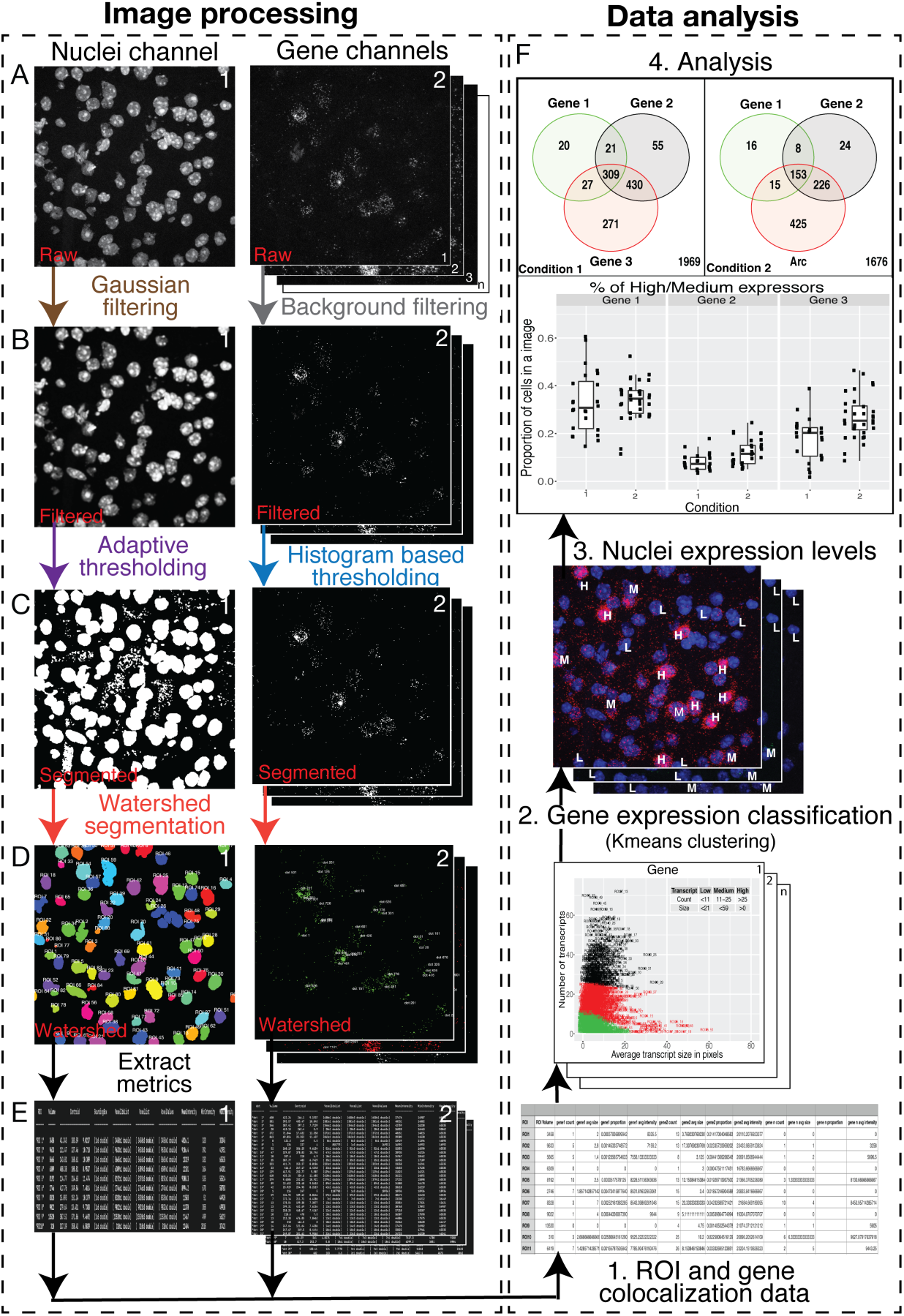
*Dotdotdot* image processing and data analysis workflows for smFISH data in mouse tissue. A-E, Image processing workflow. A) Raw *“.czi”* images of mouse nuclei (DAPI) and gene channels (i.e., *Bdnf* Ex1 [Opal520], *Arc* [Opal570], and *Bdnf* Ex4 [Opal690]). B) Processed images of nuclei and transcript channels (gaussian smoothened (B1) and background filtered (B2)). C) Segmented binary images of nuclei (adaptive thresholding) and transcript (image histogram-based thresholding) channels. D) Final watershed segmentation of nuclei and transcript channels. E) Metrics showing ID, volume (in pixels), location (centroid, bounding box, indices), intensities (mean, minimum and maximum) of each segmented region (i.e. nuclei and transcripts) per channel. F, Data analysis workflow. 1) Location metrics from E1 and E2 are used to find ROIs/nuclei and transcript colocalization information. Table shows each ROI/nucleus with its transcript quantification (count, average size, average intensity, proportion of nuclei volume occupied) per gene channel. 2-3) K-means clustering of ROIs/nuclei into high, medium and low expressers per gene channel based on transcript count and average transcript size of nuclei. 4) Data analysis based on thresholds from k-means clustering.

**Supplementary Figure 2.**
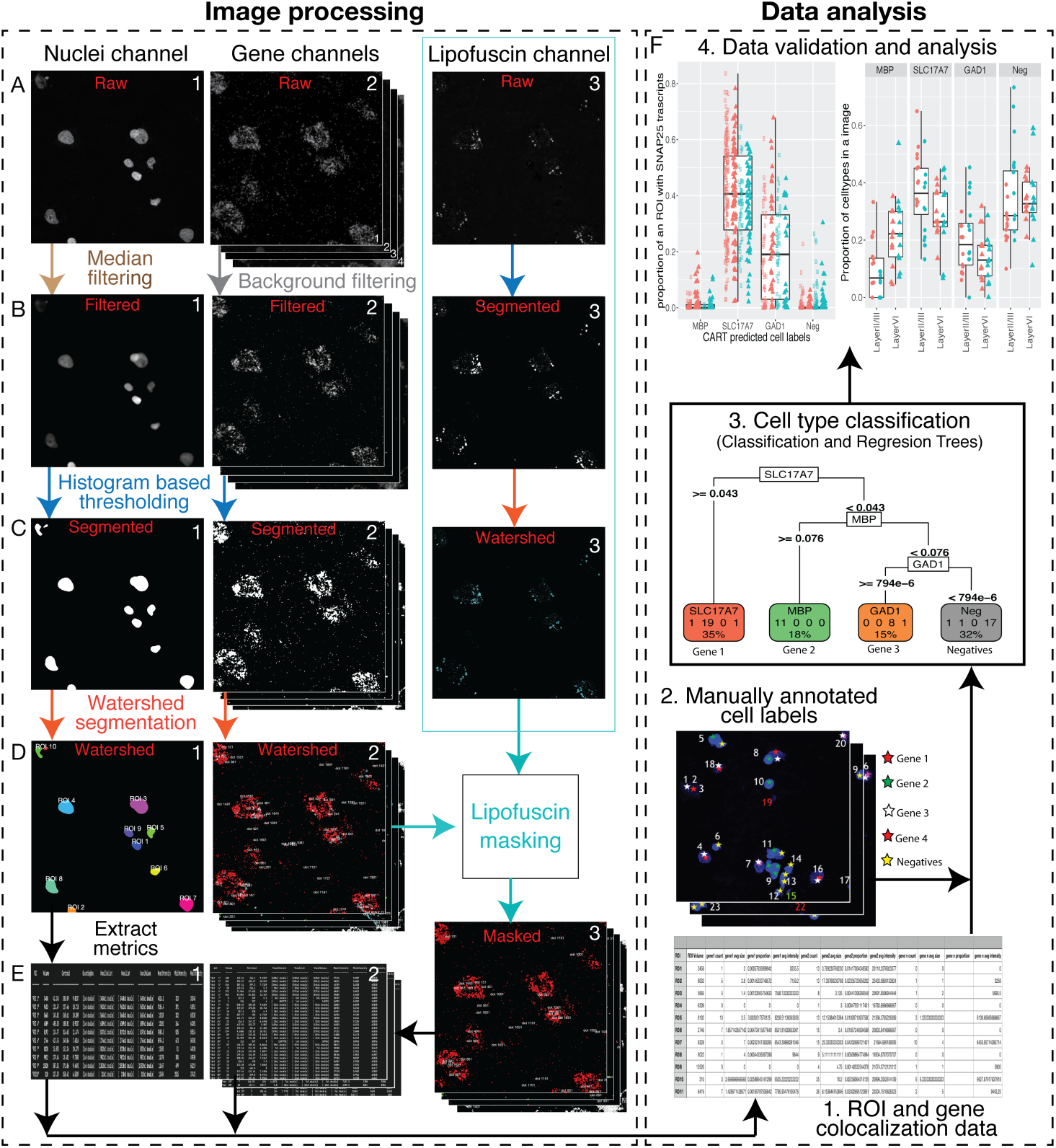
Image processing and data analysis workflows for smFISH data in human tissue. A-E, Image processing workflow. A1, A2, A3) Raw *“.czi”* images of human nuclei (DAPI), gene channels (*MBP* [Opal520], *SLC17A7* [Opal570], *GAD1* [Opal620], and *SNAP25* [Opal690]) and the lipofuscin channel. B1, B2) Processed images of nuclei and transcript channels (3D Median filtered/smoothened (B1) and background filtered (B2)). There was no processing step on the lipofuscin channel to retain all signals. C1, C2, B3) Histogram-based threshold-ed binary images of nuclei, transcript and lipofuscin channels. D1, D2, C3) Final watershed segmentation of nuclei, transcript (overlaid with lipofuscin autofluorescence pixels in cyan) and lipofuscin channels. E3) Segmented and lipofuscin-masked image of each gene channel. E1,E2) Final extracted metrics showing ID, volume (in pixels), location (centroid, bounding box, indices), intensities (mean, minimum and maximum) of each segmented region (i.e. nuclei and transcripts after lipofuscin masking) per channel. F) Data analysis workflow. 1) Location metrics from E1 and E2 are used to find ROIs/nuclei and transcript colocalization information. Table shows each ROI/nucleus with its transcript quantification (count, average size, average intensity, proportion of nuclei volume occupied) per gene channel. 2) Manually annotated images show ROIs/nuclei positive for gene targets (color labelled as red-*SLC17A7*, green-*MBP,* white-*SNAP25*, pink-*GAD1* and yellow-negative) based on qualitative analysis. 3) Classification and regression tree (CART) model produced by using the proportion of nuclei occupied by each gene and its manually annotated cell labels (Gene1 (*SLC17A7*), Gene2 (*MBP*), Gene3 (*GAD1*) and negative). 4) Based on CART predictions, nuclei from all images are classified into predefined cell types and data analysis is performed. To validate CART predictions, we show that *SNAP25* is enriched in nuclei that are positive for *SLC17A7* and *GAD1*.

**Supplementary Figure 3.**
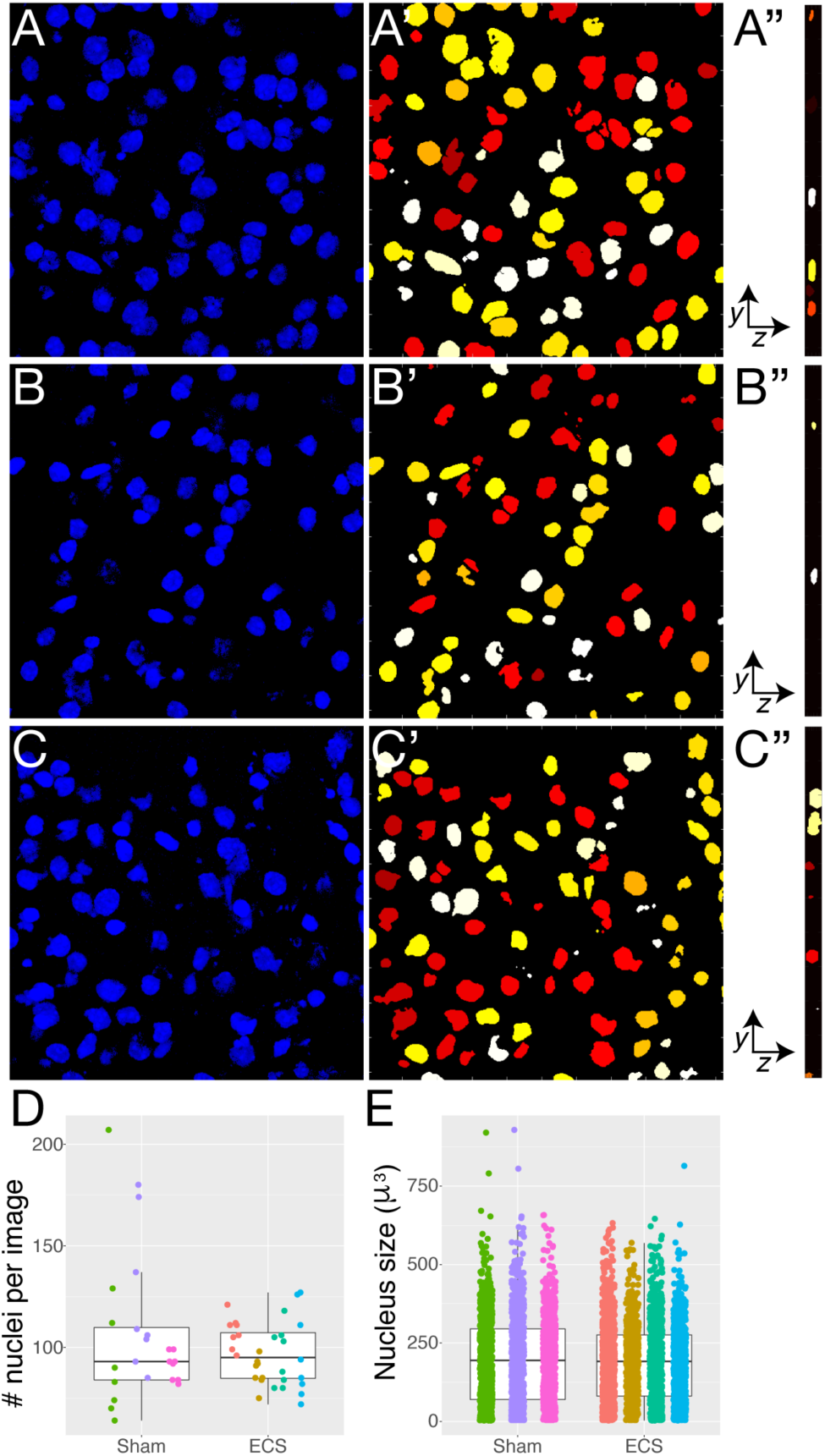
Nuclei segmentation defines regions of interest (ROIs) in mouse tissue. A-C) DAPI staining depicting individual nuclei in x,y-dimensions of a single confocal z-plane from three representative areas in the mouse piriform cortex. A’-C’) Corresponding nuclear segmentation in x,y-dimensions with each nucleus (yellow, red, orange, or white) representing a single ROI. A’’-C’’) Nuclear segmentation in y, z-dimensions. For z-stacks, nuclear segmentation is performed in each z-plane and ROIs are reconstructed in 3 dimensions. D) Number of segmented nuclei per field in images acquired from Sham and ECS treated mice (n= 24 images, 3 mice and n= 32 images, mice, respectively). Each dot represents an image and each color represents a different mouse. E) Average size of segmented nucleus in Sham compared to ECS images (n= 56 images and 5645 nuclei).

**Supplementary Figure 4.**
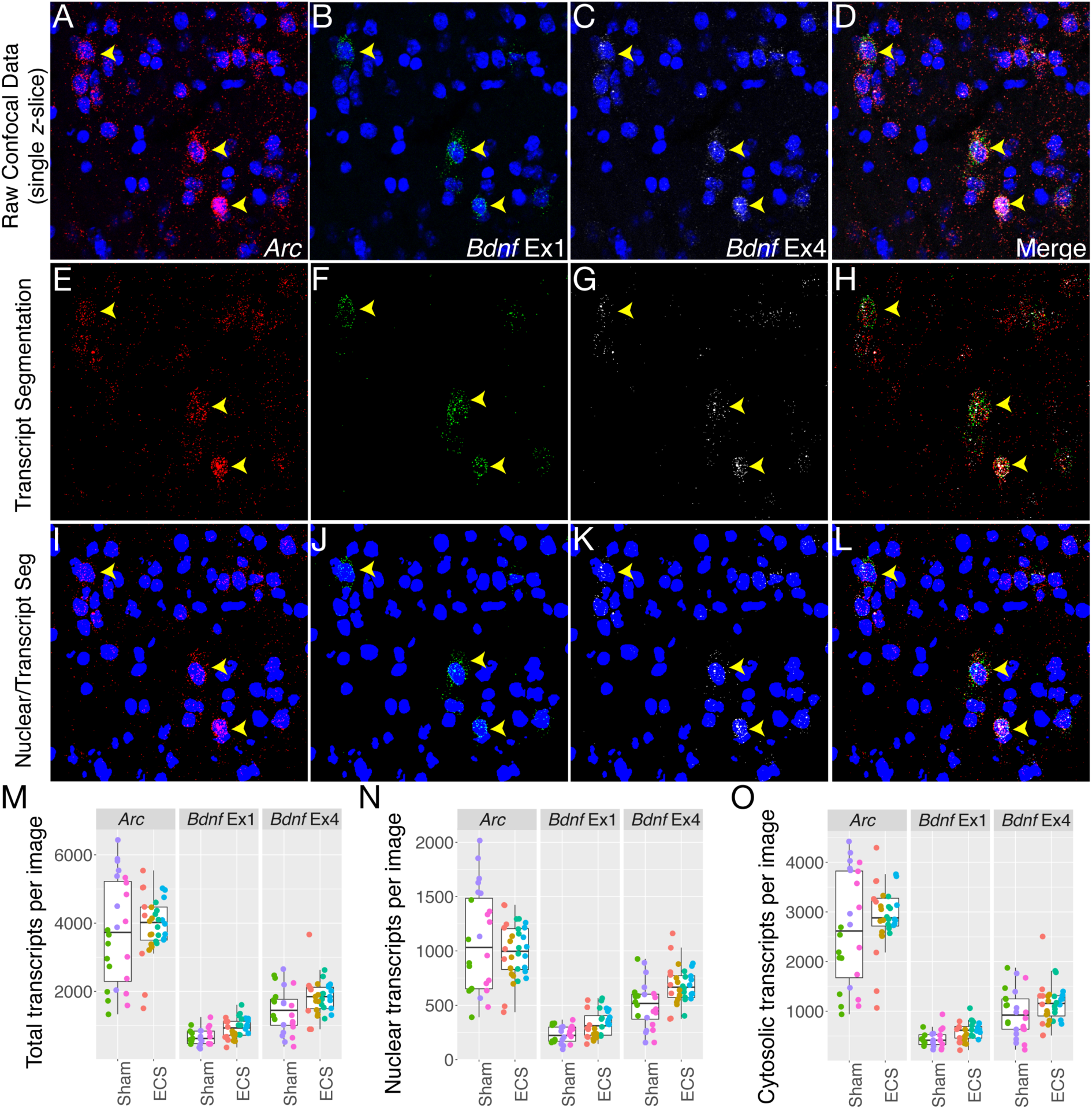
Three-dimensional dot segmentation and feature extraction delineates individual probe signals in multiplex images. A-D) Representative confocal z-plane of mouse piriform cortex depicting nuclear DAPI staining (blue) and single transcripts for *Arc* (A), *Bdnf Ex1*(B), *Bdnf Ex4* (C), and merged (D). E-H) Corresponding dot segmentation for *Arc* (E), *Bdnf Ex1* (F), *Bdnf Ex4* (G), and merged (H). I-L) Overlay of nuclear and dot segmentation used for identification of ROIs and quantification of dot/transcript features (size, number, and intensity) for *Arc* (I), *Bdnf* Ex1 (J), *Bdnf* Ex4 (K), and merged (L). M) Total number of *Arc*, *Bdnf* Ex1, and *Bdnf* Ex4 transcripts per field in images acquired from Sham and ECS treated mice (n= 3 mice, 24 images, 2543 nuclei and n=4 mice, 32 images, and 3102nuclei, respectively). Each dot represents an image and each color represents a different mouse. N) Total number of nuclear transcripts per field for *Arc*, *Bdnf* Ex1, and *Bdnf* Ex4 in Sham compared to ECS. O) Total number of cytosolic transcripts per field for *Arc*, *Bdnf* Ex1, and *Bdnf* Ex4 in Sham compared to ECS. Yellow arrows depict positive cells.

**Supplementary Figure 5.**
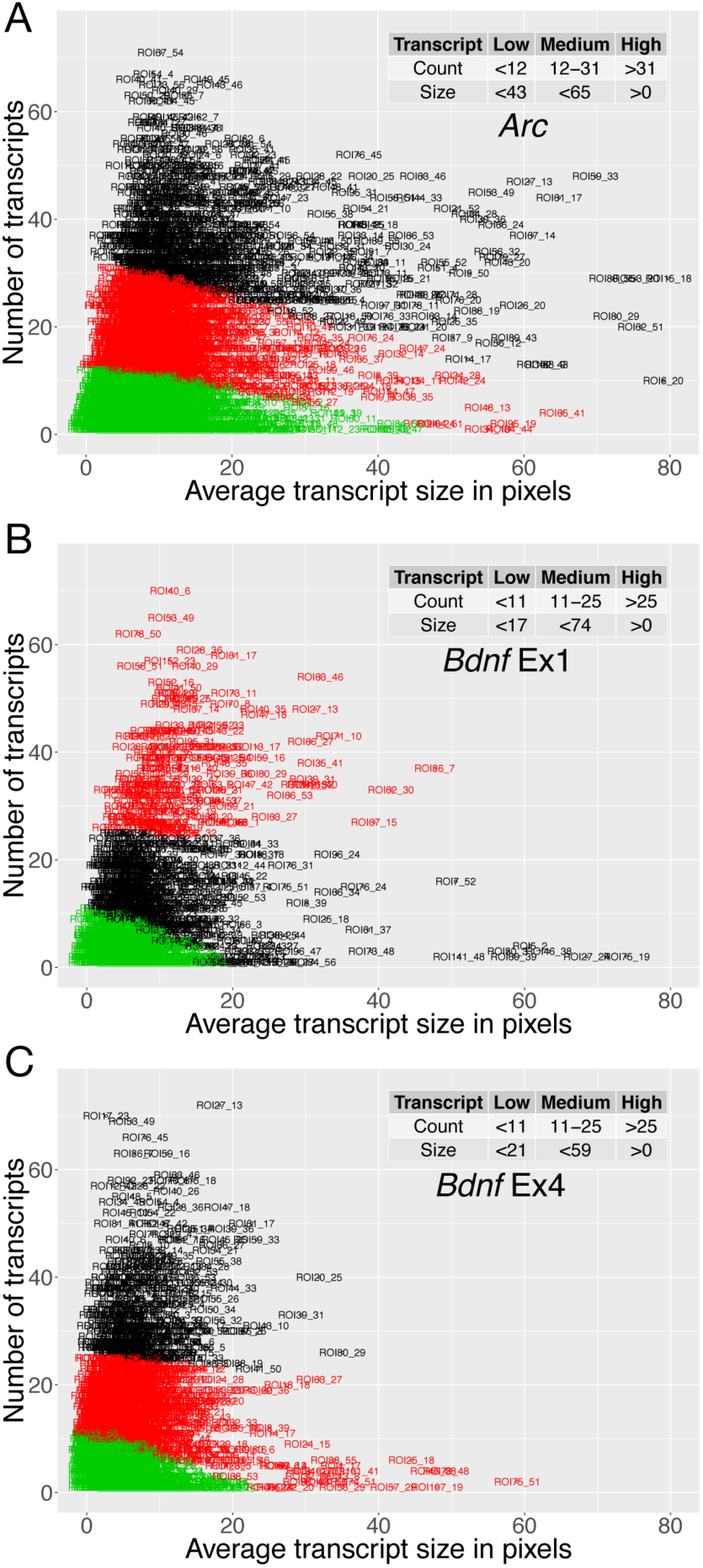
K-means cluster analysis uses transcript (dot) count and size per ROI to group ROIs as low, medium, or high expressers for individual ARGs. A) Plot depicting clustering of 4,967 ROIs with low, medium, and high *Arc* expression based on number of transcripts (low: <12; medium: 12-31; high: >31) and average transcript size (low: <43; medium: <65; high: >0). B) Plot depicting clustering of 2,538 ROIs with low, medium, and high *Bdnf* Ex1 expression based on number of transcripts (low: <11; medium: 11-25; high: >25) and average transcript size (low: <17; medium: <74; high: >0). C) Plot depicting clustering of 3,983 ROIs with low, medium, and high *Bdnf* Ex4 expression based on number of transcripts (low: <11; medium: 11-25; high: >25) and average transcript size (low: <21; medium: <59; high: >0). Nuclei with <1 transcript are directly assigned to the low group.

**Supplementary Figure 6.**
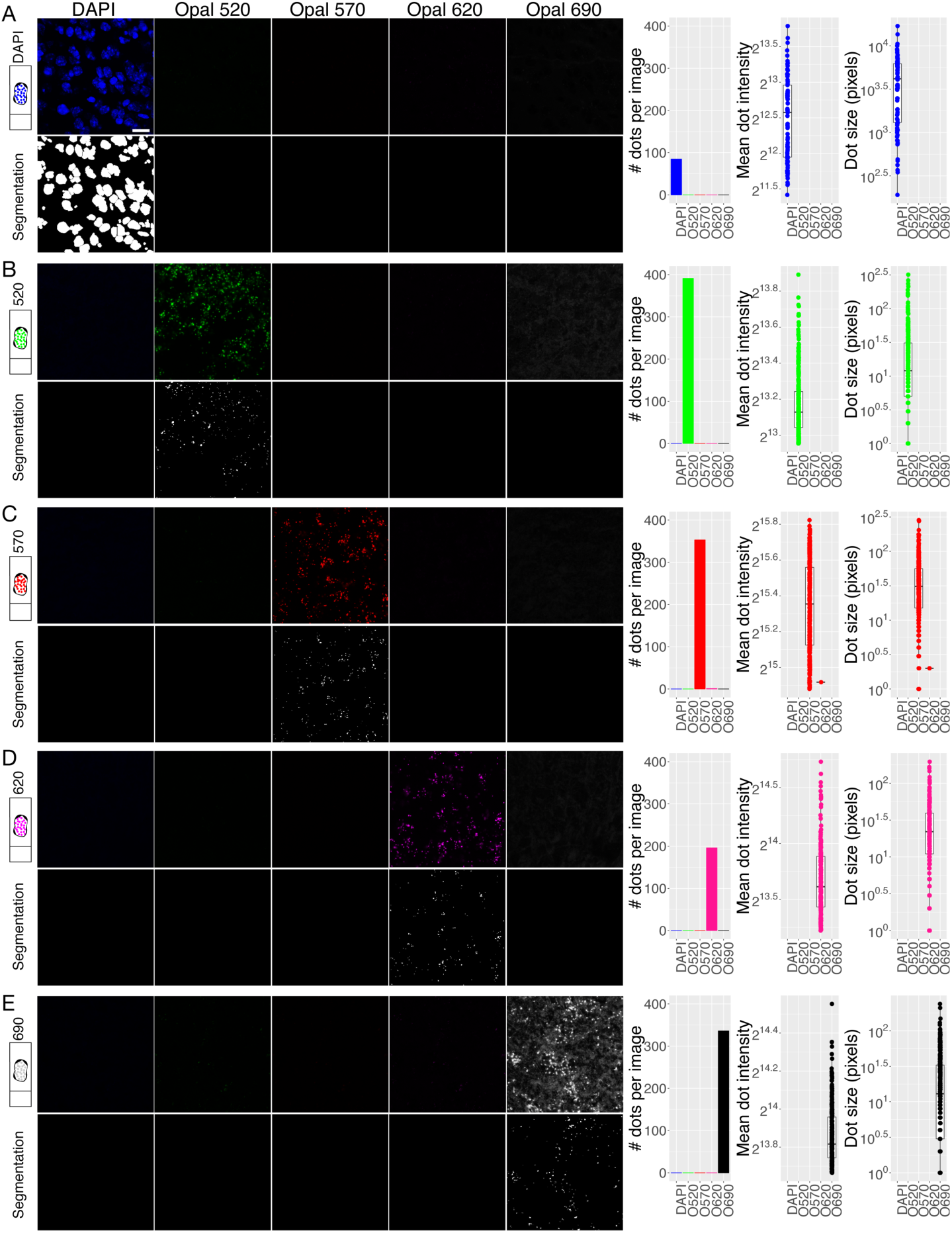
Validation of reference emission spectral profiles (“fingerprints”) used for linear unmixing. A) A reference emission spectral profile, or “fingerprint,” was created in Zen software for DAPI using mouse brain tissue subjected to pretreatment conditions, but no additional probe labeling. Top panel shows linear unmixing with the DAPI fingerprint specifically recognizes DAPI signal. Bottom panel depicts segmentation for ROI identification. B-E) Reference emission spectral profiles were created in Zen software for each of the Opal dyes (referred to as Opal (O)520, Opal (O)570, Opal (O)620, and Opal(O) 690) using a “single positive” slide of mouse brain tissue hybridized with a positive control probe against a house-keeping gene, *POLR2A*, and visualized with the respective Opal dye (Opal520 in B, Opal570 in C, Opal620 in D, and Opal690 in E). Top panels show linear unmixing with DAPI, Opal520, Opal570, Opal620, and Opal690 for each single positive slide. Bottom panels depict segmentation for dot detection. Plots to right of images show *POLR2A* dot number, size, and average intensity after linear unmixing of respective single positive images. Importantly, reference emission spectral profiles are highly specific (i.e. when *POLR2A* is labeled with Opal570 in C, the spectrally similar Opal620 fingerprint does not detect Opal570 signal, etc.). Similarly, dot features are only detected after unmixing with the relevant reference emission spectral profile. Scale bar is 20um.

**Supplementary Figure 7.**
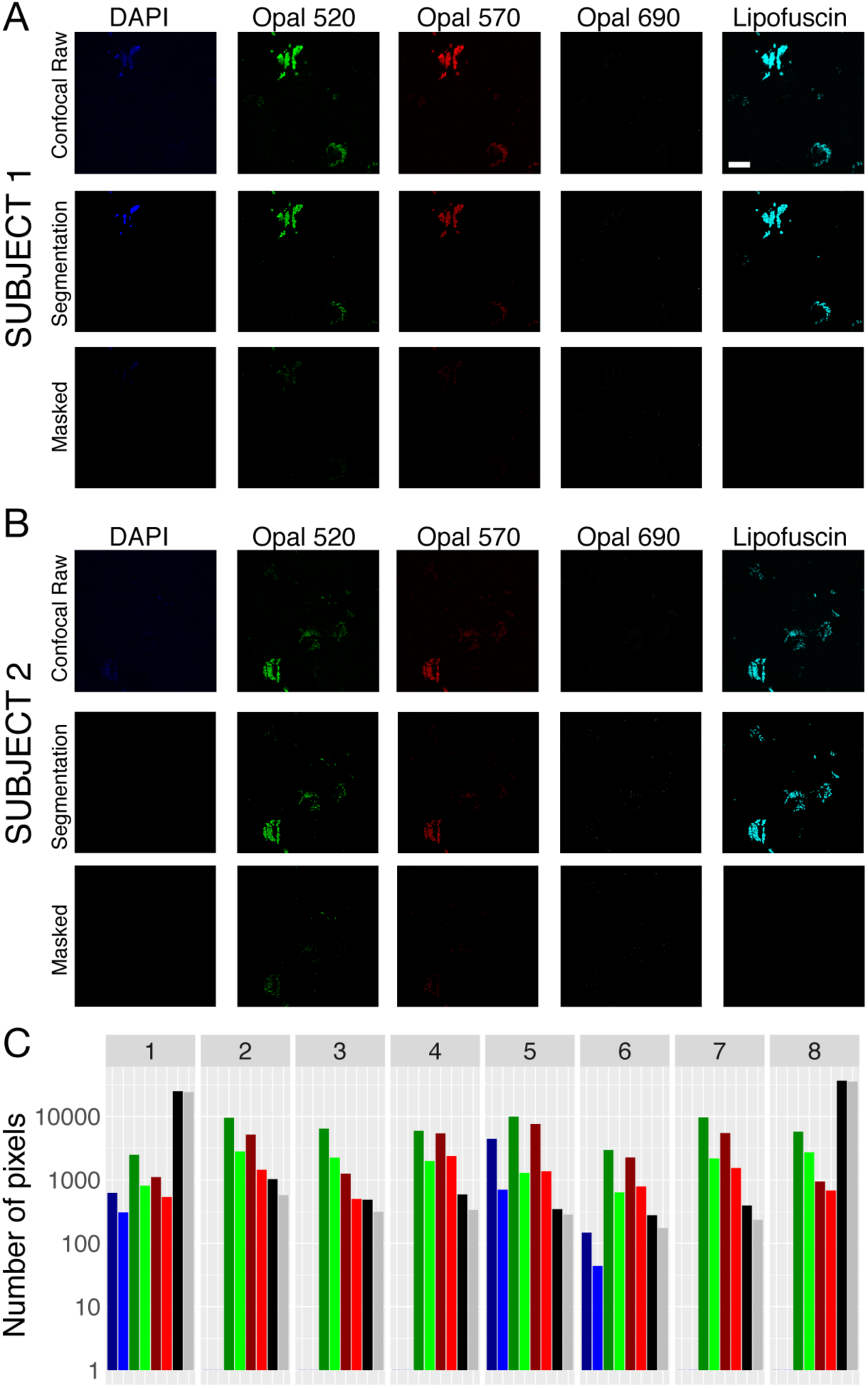
A common spectral signature for lipofuscin can be used across multiple subjects to identify and exclude autofluorescence. A-B) A reference emission spectral profile, or “fingerprint,” was created in Zen software for lipofuscin autofluorescence using DLPFC from a representative subject hybridized with a negative control probe for the bacterial gene *dapB*. For 2 different human subjects (A and B), top panels show a single confocal z-slice after linear unmixing with DAPI, Opal520, Opal570, Opal60, and lipofuscin fingerprints. Middle panels show corresponding segmentation. Bottom panels show segmentation after masking with lipofuscin signals. C) Plot depicting the number of transcript pixels detected for DAPI (blue bars), Opal520 (green bars), Opal570 (red bars), Opal690 (black/gray bars) before and after masking with lipofuscin (dark bars=before, light bars=after) across 8 images derived from 4 different subjects (Images 5-6 from Subject 1, Images 7-8 from Subject 2, Images 1-2 from Subject 3, Images 3-4 from Subject 4,). The reference emission spectral profile for lipofuscin is equally effective across subjects and successfully masks and excludes pixels confounded by autofluorescence across the electromagnetic spectrum. Scale bar 20um.

**Supplementary Figure 8.**
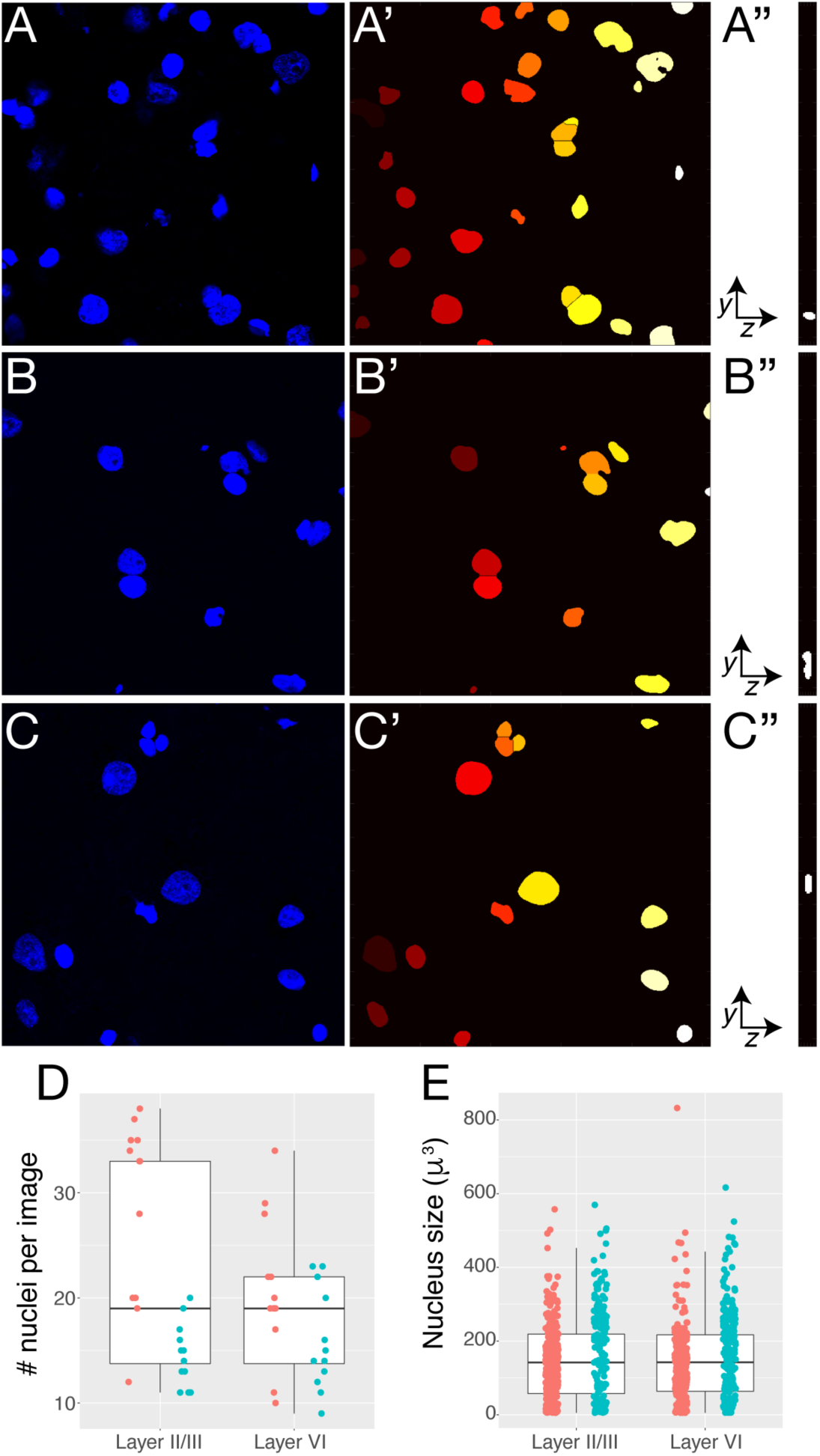
Nuclei segmentation defines regions of interest (ROIs) in human brain. A-C) DAPI staining depicting individual nuclei in *x*,*y*-dimensions of a single confocal *z*-plane from three representative areas in postmortem human DLPFC. A’-C’) Corresponding nuclear segmentation in *x*,*y*-dimensions with each nucleus (yellow, red, orange, or white) representing a single ROI. A’’-C’’) Nuclear segmentation in *y*, *z*-dimensions. For *z*-stacks, nuclear segmentation is performed in each z-plane and ROIs are reconstructed in 3 dimensions. D) Number of segmented nuclei per field in images acquired from cortical layers II/III and VI (n=2 subjects, 2 cortical strips per subject, 24 images). E) Average size of segmented nucleus in layer II/III compared to layer VI (n= 519 and 442 nuclei, respectively).

**Supplementary Figure 9.**
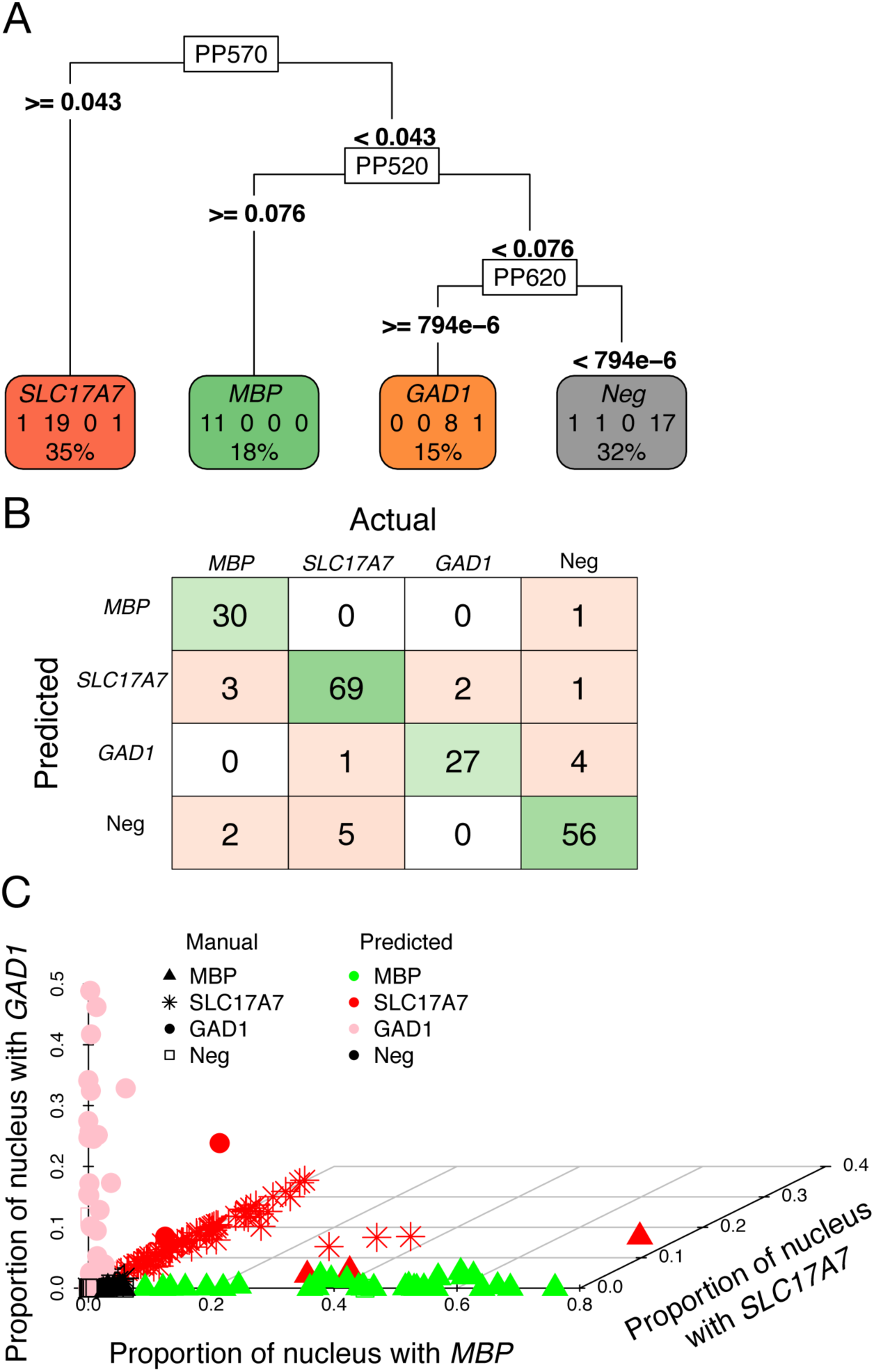
Classification and regression tree (CART) model for accurately predicting cell types in DLPFC. A) Classification tree built from 60 random ROIs from 5 manually annotated images using the rpart algorithm with termination criteria of 4 major classes: *SLC17A7* (excitatory neuron), *MBP* (oligodendrocyte), *GAD1* (inhibitory neuron), and triple negative (Neg; likely microglia and astrocytes). For example, a cell is defined as an *SLC17A7*+ excitatory neuron when the volume of the ROI is at least 4.3% *SLC17A7* (given that *MBP* covers less than 7.6% of the ROI and *GAD1* covers less than 0.0794% of the ROI). For these 60 ROIs, the classifier predicts 35% of the ROIs are SLC17A7, 18% are MBP, 15% are GAD1, and 32% are triple negative. B) Confusion matrix for 201 manually annotated ROIs comparing predicted and actual cell type. C) Plot showing manual and predicted cell type for each ROI plotted against the proportion of the nuclei positive for *GAD1*, *SLC17A7*, and *MBP*. Prediction accuracy of the CART model is 91%.

**Supplementary Figure 10.**
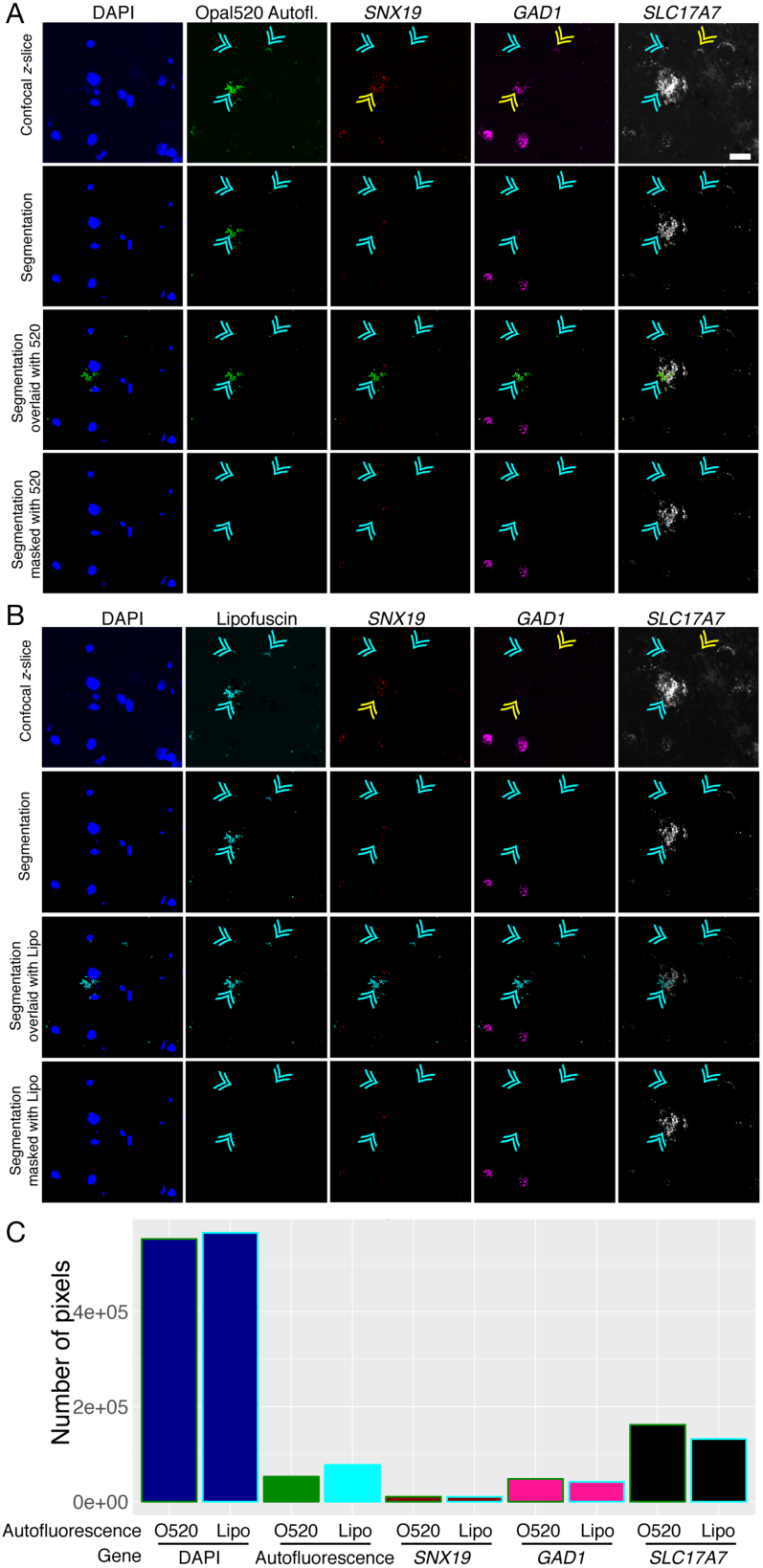
Flexibility for using autofluorescence detected in a narrow spectral range to mask lipofuscin autofluorescence. A) Top panel shows confocal z-slice after linear unmixing with DAPI, Opal520, Opal570, Opal620, and Opal690 fingerprints to detect *SNX19* (Opal570), *GAD1* (Opal620), and *SLC17A7* (Opal690). As no probe was used with the Opal520 dye, signal in this spectral range can be attributed to lipofuscin autofluorescence. Second panel shows segmentation of fluorescent signals. Third panel shows segmentation overlaid with autofluorescence captured by unmixing with the Opal520 fingerprint. Bottom panel shows segmentation for each gene after masking with Opal520 autofluorescence. Cyan and yellow arrows show lipofuscin autofluorescence. B) Using the same lambda stack, top panel shows confocal z-slice after linear unmixing with DAPI, lipofuscin, Opal570, Opal620, and Opal690 fingerprints to detect *SNX19* (Opal570), *GAD1* (Opal620), *SLC17A7* (Opal690), and lipofuscin autofluorescence. Second panel shows segmentation of fluorescent signals. Third panel shows segmentation overlaid with autofluorescence captured by unmixing with the lipofuscin fingerprint. Bottom panel shows segmentation for each gene after masking with lipofuscin autofluorescence. Lipofuscin fingerprint captures additional autofluorescence compared to Opal520 fingerprint (compare yellow arrows in A and B). C) Plot comparing the number of pixels detected for DAPI or each gene when lipofuscin autofluorescence is unmixed using the Opal520 (O520; green border bars) versus lipofuscin (Lipo; cyan border bars) fingerprints. The lipofuscin fingerprint captures more autofluorescent pixels than the Opal520 fingerprint (cyan vs. green bar) thereby reducing the number of pixels attributed to autofluorescence in other channels. Scale bar is 20um.

**Supplementary Figure 11.**
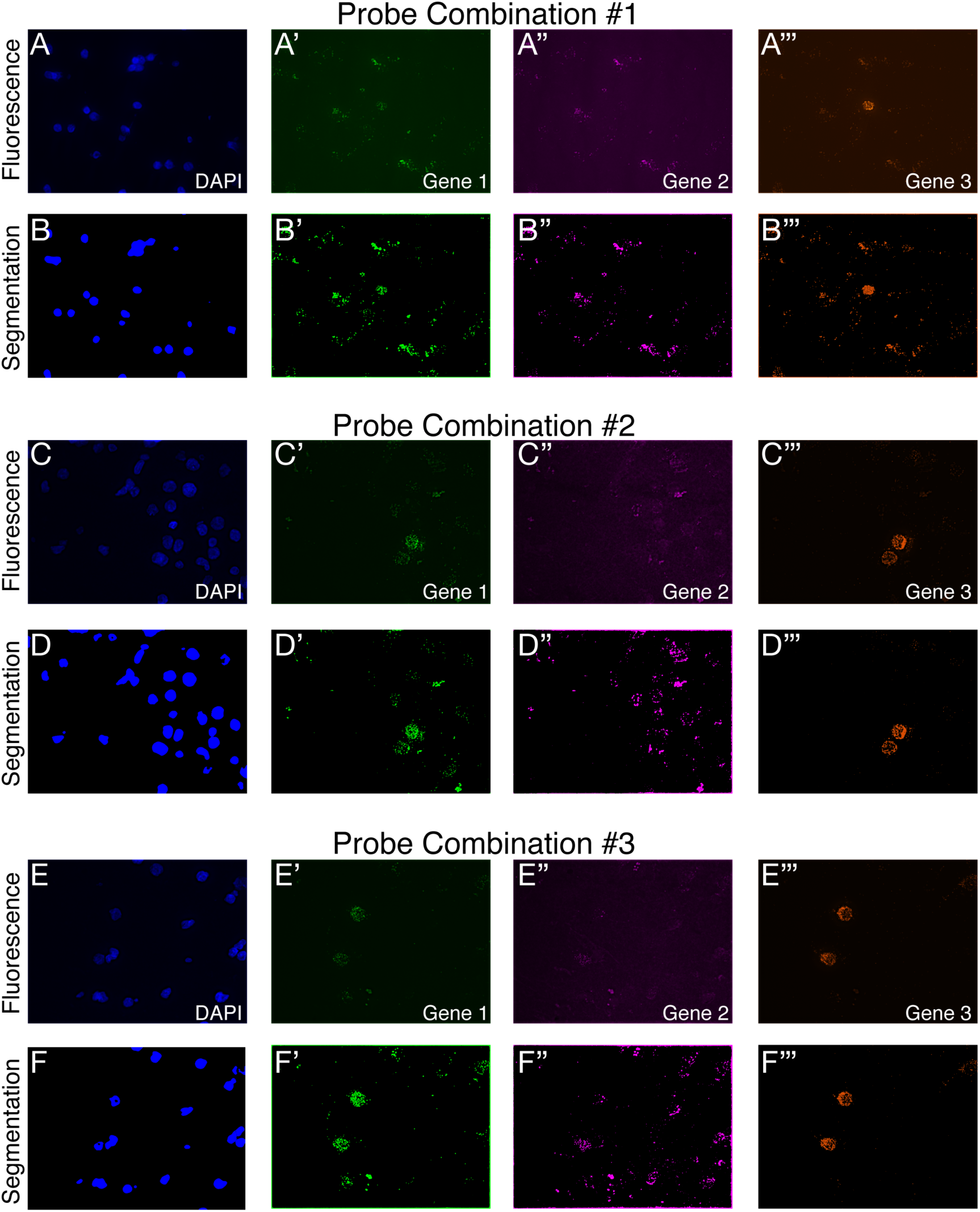
*dotdotdot* is compatible with diverse file formats generated using different microscope systems. A-F) Raw confocal fluorescence (A, C, E) and corresponding nuclear/transcript segmentation (B, D, F) from 3 unique RNAscope images acquired in postmortem human brain tissue utilizing separate probe combinations. Imaging processing with *dotdotdot* was performed on Nikon “.*nd2*” files.

## Notes

https://github.com/LieberInstitute/dotdotdot

## REFERENCES

1. Poulin, J.-F., Tasic, B., Hjerling-Leffler, J., Trimarchi, J. M. & Awatramani, R. Disentangling neural cell diversity using single-cell transcriptomics. Nat. Neurosci. 19, 1131–1141 (2016).

2. Lein, E., Borm, L. E. & Linnarsson, S. The promise of spatial transcriptomics for neuroscience in the era of molecular cell typing. Science 358, 64–69 (2017).

3. Lake, B. B. et al. Neuronal subtypes and diversity revealed by single-nucleus RNA sequencing of the human brain. Science 352, 1586–1590 (2016).

4. Hodge, R. D. et al. Conserved cell types with divergent features in human versus mouse cortex. Nature (2019). doi:10.1038/s41586-019-1506-7

5. Boldog, E. et al. Transcriptomic and morphophysiological evidence for a specialized human cortical GABAergic cell type. Nat. Neurosci. 21, 1185–1195 (2018).

6. Tartt, A. N. et al. Considerations for assessing the extent of hippocampal neurogenesis in the adult and aging human brain. Cell Stem Cell 23, 782–783 (2018).

7. Sorrells, S. F. et al. Human hippocampal neurogenesis drops sharply in children to undetectable levels in adults. Nature 555, 377–381 (2018).

8. McNicol, A. M. & Farquharson, M. A. In situ hybridization and its diagnostic applications in pathology. J. Pathol. 182, 250–261 (1997).

9. Weickert, C. S., Rothmond, D. A. & Purves-Tyson, T. D. Considerations for optimal use of postmortem human brains for molecular psychiatry: lessons from schizophrenia. Handb. Clin. Neurol. 150, 221–235 (2018).

10. Wang, F. et al. RNAscope: a novel in situ RNA analysis platform for formalin-fixed, paraffin-embedded tissues. J. Mol. Diagn. 14, 22–29 (2012).

11. Wang, Z. et al. Automated quantitative RNA in situ hybridization for resolution of equivocal and heterogeneous ERBB2 (HER2) status in invasive breast carcinoma. J. Mol. Diagn. 15, 210–219 (2013).

12. Tsanov, N. et al. smiFISH and FISH-quant - a flexible single RNA detection approach with super-resolution capability. Nucleic Acids Res. 44, e165 (2016).

13. Dowson, J. H. & Harris, S. J. Quantitative studies of the autofluorescence derived from neuronal lipofuscin. J. Microsc. 123, 249–258 (1981).

14. Benavides, S. H., Monserrat, A. J., Fariña, S. & Porta, E. A. Sequential histochemical studies of neuronal lipofuscin in human cerebral cortex from the first to the ninth decade of life. Arch. Gerontol. Geriatr. 34, 219–231 (2002).

15. Baleriola, J. et al. Axonally synthesized ATF4 transmits a neurodegenerative signal across brain regions. Cell 158, 1159–1172 (2014).

16. Bissel, S. J. et al. Human parechovirus 3 meningitis and fatal leukoencephalopathy. J. Neuropathol. Exp. Neurol. 74, 767–777 (2015).

17. Bialas, A. R. et al. Microglia-dependent synapse loss in type I interferon-mediated lupus. Nature 546, 539–543 (2017).

18. Gandal, M. J. et al. Transcriptome-wide isoform-level dysregulation in ASD, schizophrenia, and bipolar disorder. Science 362, (2018).

19. Smith, R. S. et al. Sodium channel SCN3A (nav1.3) regulation of human cerebral cortical folding and oral motor development. Neuron 99, 905–913.e7 (2018).

20. Fish, K. N., Rocco, B. R. & Lewis, D. A. Laminar distribution of subsets of gabaergic axon terminals in human prefrontal cortex. Front. Neuroanat. 12, 9 (2018).

21. Rocco, B. R., Oh, H., Shukla, R., Mechawar, N. & Sibille, E. Fluorescence-based cell-specific detection for laser-capture microdissection in human brain. Sci. Rep. 7, 14213 (2017).

22. Jolly, S. et al. Single-Cell Quantification of mRNA Expression in The Human Brain. Sci. Rep. 9, 12353 (2019).

23. Pyon, W. S., Gray, D. T. & Barnes, C. A. An Alternative to Dye-Based Approaches to Remove Background Autofluorescence From Primate Brain Tissue. Front. Neuroanat. 13, 73 (2019).

24. Shah, S., Lubeck, E., Zhou, W. & Cai, L. seqFISH Accurately Detects Transcripts in Single Cells and Reveals Robust Spatial Organization in the Hippocampus. Neuron 94, 752–758.e1 (2017).

25. Chen, K. H., Boettiger, A. N., Moffitt, J. R., Wang, S. & Zhuang, X. RNA imaging. Spatially resolved, highly multiplexed RNA profiling in single cells. Science 348, aaa6090 (2015).

26. Ke, R. et al. In situ sequencing for RNA analysis in preserved tissue and cells. Nat. Methods 10, 857–860 (2013).

27. Lee, J. H. et al. Highly multiplexed subcellular RNA sequencing in situ. Science 343, 1360–1363 (2014).

28. Wang, X. et al. Three-dimensional intact-tissue sequencing of single-cell transcriptional states. Science 361, (2018).

29. Altar, C. A. et al. Electroconvulsive seizures regulate gene expression of distinct neurotrophic signaling pathways. J. Neurosci. 24, 2667–2677 (2004).

30. Timmusk, T. et al. Multiple promoters direct tissue-specific expression of the rat BDNF gene. Neuron 10, 475–489 (1993).

31. Aid, T., Kazantseva, A., Piirsoo, M., Palm, K. & Timmusk, T. Mouse and rat BDNF gene structure and expression revisited. J. Neurosci. Res. 85, 525–535 (2007).

32. Colliva, A., Maynard, K. R., Martinowich, K. & Tongiorgi, E. in Brain-Derived Neurotrophic Factor (BDNF) (eds. Duarte, C. B. & Tongiorgi, E.) 143, 27–53 (Springer New York, 2019).

33. Maynard, K. R. et al. Functional Role of BDNF Production from Unique Promoters in Aggression and Serotonin Signaling. Neuropsychopharmacology 41, 1943–1955 (2016).

34. Hong, E. J., McCord, A. E. & Greenberg, M. E. A biological function for the neuronal activity-dependent component of Bdnf transcription in the development of cortical inhibition. Neuron 60, 610–624 (2008).

35. Lyford, G. L. et al. Arc, a growth factor and activity-regulated gene, encodes a novel cytoskeleton-associated protein that is enriched in neuronal dendrites. Neuron 14, 433–445 (1995).

36. Ma, L. et al. Schizophrenia risk variants influence multiple classes of transcripts of sorting nexin 19 (SNX19). Mol. Psychiatry (2019). doi:10.1038/s41380-018-0293-0

37. Larsen, M. H. et al. Regulation of activity-regulated cytoskeleton protein (Arc) mRNA after acute and chronic electroconvulsive stimulation in the rat. Brain Res. 1064, 161–165 (2005).

38. Vismer, M. S., Forcelli, P. A., Skopin, M. D., Gale, K. & Koubeissi, M. Z. The piriform, perirhinal, and entorhinal cortex in seizure generation. Front. Neural Circuits 9, 27 (2015).

39. Revuelta, M., Castaño, A., Venero, J. L., Machado, A. & Cano, J. Long-lasting induction of brain-derived neurotrophic factor is restricted to resistant cell populations in an animal model of status epilepticus. Neuroscience 103, 955–969 (2001).

40. Sun, W. et al. Identification of novel electroconvulsive shock-induced and activity-dependent genes in the rat brain. Biochem. Biophys. Res. Commun. 327, 848–856 (2005).

41. Maynard, K. R., Hobbs, J. W., Rajpurohit, S. K. & Martinowich, K. Electroconvulsive seizures influence dendritic spine morphology and BDNF expression in a neuroendocrine model of depression. Brain Stimulat. 11, 856–859 (2018).

42. DeNardo, L. A. et al. Temporal evolution of cortical ensembles promoting remote memory retrieval. Nat. Neurosci. 22, 460–469 (2019).

43. Denny, C. A. et al. Hippocampal memory traces are differentially modulated by experience, time, and adult neurogenesis. Neuron 83, 189–201 (2014).

44. West, A. E., Pruunsild, P. & Timmusk, T. Neurotrophins: transcription and translation. Handb. Exp. Pharmacol. 220, 67–100 (2014).

45. Baj, G., Leone, E., Chao, M. V. & Tongiorgi, E. Spatial segregation of BDNF transcripts enables BDNF to differentially shape distinct dendritic compartments. Proc Natl Acad Sci USA 108, 16813–16818 (2011).

46. Pattabiraman, P. P. et al. Neuronal activity regulates the developmental expression and subcellular localization of cortical BDNF mRNA isoforms in vivo. Mol. Cell. Neurosci. 28, 556–570 (2005).

47. Molyneaux, B. J., Arlotta, P., Menezes, J. R. L. & Macklis, J. D. Neuronal subtype specification in the cerebral cortex. Nat. Rev. Neurosci. 8, 427–437 (2007).

48. Sun, Y., Ip, P. & Chakrabartty, A. Simple Elimination of Background Fluorescence in Formalin-Fixed Human Brain Tissue for Immunofluorescence Microscopy. J. Vis. Exp. (2017). doi:10.3791/56188

49. Moffitt, J. R. et al. High-performance multiplexed fluorescence in situ hybridization in culture and tissue with matrix imprinting and clearing. Proc Natl Acad Sci USA 113, 14456–14461 (2016).

50. Sylwestrak, E. L., Rajasethupathy, P., Wright, M. A., Jaffe, A. & Deisseroth, K. Multiplexed Intact-Tissue Transcriptional Analysis at Cellular Resolution. Cell 164, 792–804 (2016).

51. Guintivano, J., Aryee, M. J. & Kaminsky, Z. A. A cell epigenotype specific model for the correction of brain cellular heterogeneity bias and its application to age, brain region and major depression. Epigenetics 8, 290–302 (2013).

52. Hallock, H. L. et al. Manipulation of a genetically and spatially defined sub-population of BDNF-expressing neurons potentiates learned fear and decreases hippocampal-prefrontal synchrony in mice. Neuropsychopharmacology (2019). doi:10.1038/s41386-019-0429-1

53. MacQueen, J. Some methods for classification and analysis of multivariate observations. (1967).

54. Breiman, L., Friedman, J. H., Olshen, R. A. & Stone, C. J. Classification and regression trees. (Wadsworth & Brooks/Cole Advanced Books & Software, 1984). doi:10.1201/9781315139470

55. Alegro, M. et al. Automating cell detection and classification in human brain fluorescent microscopy images using dictionary learning and sparse coding. J. Neurosci. Methods 282, 20–33 (2017).

56. Strell, C. et al. Placing RNA in context and space - methods for spatially resolved transcriptomics. FEBS J. 286, 1468–1481 (2019).

57. Burgess, D. J. Spatial transcriptomics coming of age. Nat. Rev. Genet. 20, 317 (2019).

58. Maynard, K. R., Jaffe, A. E. & Martinowich, K. Spatial transcriptomics: putting genome-wide expression on the map. Neuropsychopharmacology (2019). doi:10.1038/s41386-019-0484-7

59. Ståhl, P. L. et al. Visualization and analysis of gene expression in tissue sections by spatial transcriptomics. Science 353, 78–82 (2016).

60. Rodriques, S. G. et al. Slide-seq: A scalable technology for measuring genome-wide expression at high spatial resolution. Science 363, 1463–1467 (2019).

61. Svensson, V., Teichmann, S. A. & Stegle, O. SpatialDE: identification of spatially variable genes. Nat. Methods 15, 343–346 (2018).

62. Edsgärd, D., Johnsson, P. & Sandberg, R. Identification of spatial expression trends in single-cell gene expression data. Nat. Methods 15, 339–342 (2018).

63. Maniatis, S. et al. Spatiotemporal dynamics of molecular pathology in amyotrophic lateral sclerosis. Science 364, 89–93 (2019).

64. Moncada, R. et al. Building a tumor atlas: integrating single-cell RNA-Seq data with spatial transcriptomics in pancreatic ductal adenocarcinoma. BioRxiv (2018). doi:10.1101/254375

65. Schloesser, R. J. et al. Antidepressant-like Effects of Electroconvulsive Seizures Require Adult Neurogenesis in a Neuroendocrine Model of Depression. Brain Stimulat. 8, 862–867 (2015).

66. Lipska, B. K. et al. Critical factors in gene expression in postmortem human brain: Focus on studies in schizophrenia. Biol. Psychiatry 60, 650–658 (2006).

67. matrix laboratory, C. B. M. Matlab. (MathWorks, 2018). at <https://www.mathworks.com/products/matlab.html>

68. OME, B.-F. bfmatlab - Using Bio-Formats in MATLAB. (University of Dundee & Open Microscopy Environment, 2017). at <https://docs.openmicroscopy.org/bio-formats/5.7.3/developers/matlab-dev.html>

69. Hodneland, E., Kögel, T., Frei, D. M., Gerdes, H.-H. & Lundervold, A. CellSegm - a MATLAB toolbox for high-throughput 3D cell segmentation. Source Code Biol. Med. 8, 16 (2013).

70. Ram, S., Rodríguez, J. J. & Bosco, G. Segmentation and detection of fluorescent 3D spots. Cytometry A 81, 198–212 (2012).

71. Team, R. C. R: A Language and Environment for Statistical Computing. (2017).

72. Tests in Linear Mixed Effects Models [R package lmerTest version 3.1-0]. at <https://cran.r-project.org/web/packages/lmerTest/index.html>

